# Residues 2-7 of α-synuclein regulate amyloid formation via lipid-dependent and -independent pathways

**DOI:** 10.1101/2024.05.24.595537

**Authors:** Katherine M. Dewison, Benjamin Rowlinson, Jonathan M. Machin, Joel A. Crossley, Dev Thacker, Martin Wilkinson, Sabine M. Ulamec, G. Nasir Khan, Neil A. Ranson, Patricija van Oosten-Hawle, David J. Brockwell, Sheena E. Radford

**Affiliations:** Astbury Centre for Structural Molecular Biology, School of Molecular and Cellular Biology, Faculty of Biological Sciences, University of Leeds, Leeds LS2 9JT, United Kingdom; Department of Biological Sciences, University of North Carolina at Charlotte, Charlotte, NC, USA

**Author notes:** Corresponding authors: Sheena E. Radford; David J. Brockwell.

**Keywords:** alpha-synuclein, aggregation, amyloid, liposome, membrane destabilisation

## Abstract

Amyloid formation by α-synuclein (αSyn) occurs in Parkinson’s disease, multiple system atrophy, and dementia with Lewy bodies. Deciphering the residues that regulate αSyn amyloid fibril formation will not only provide mechanistic insight, but may also reveal new targets to prevent and treat disease. Previous investigations have identified several regions of αSyn to be important in the regulation of amyloid formation, including the non-amyloid-β component (NAC), P1 region (residues 36-42), and residues in the C-terminal domain. Recent studies have also indicated the importance of the N-terminal region of αSyn for both its physiological and pathological roles. Here, the role of residues 2-7 in the N-terminal region of αSyn are investigated in terms of their ability to regulate amyloid fibril formation *in vitro* and *in vivo*. Deletion of these residues (αSynΔN7) slows the rate of fibril formation *in vitro* and reduces the capacity of the protein to be recruited by wild-type (αSynWT) fibril seeds, despite cryo-EM showing a fibril structure consistent with those of full-length αSyn. Strikingly, fibril formation of αSynΔN7 is not induced by liposomes, despite the protein binding to liposomes with similar affinity to αSynWT. A *Caenorhabditis elegans* model also showed that αSynΔN7::YFP forms few puncta and lacks motility and lifespan defects typified by expression of αSynWT::YFP. Together, the results demonstrate the involvement of residues 2-7 of αSyn in amyloid formation, revealing a new target for the design of amyloid inhibitors that may leave the functional role of the protein in membrane binding unperturbed.

**Significance Statement:** Amyloid formation of α-synuclein (αSyn) is associated with Parkinson’s disease. Attempts to target αSyn aggregation to treat synucleinopathies, thus far, have been unsuccessful. A better understanding of residues that regulate amyloid formation may reveal new targets for therapeutics. Here, six residues at the N-terminus of αSyn are identified as regulators of amyloid formation. Deletion of these residues slows lipid-independent assembly, ablates lipid-dependent amyloid formation *in vitro*, and prevents aggregation and its associated cellular toxicity *in vivo*. Importantly, these residues are not necessary for binding to synthetic membranes. The work reveals a new target for the prevention of synucleinopathies by disfavouring aggregation without perturbing membrane binding, a property considered to be essential for the physiological function of αSyn at the synapse.

## Introduction

α-Synuclein (αSyn) is an intrinsically disordered protein that aggregates and forms amyloid in Parkinson’s disease, multiple system atrophy and dementia with Lewy bodies (1). Although these synucleinopathies can be caused by familial mutations in the *SNCA* gene encoding αSyn, such as A30P/G (2), E46K (3), H50Q (4), G51D (5), and A53T/V/E (6), the majority of cases are sporadic (7). The triggers of sporadic synucleinopathies remain unclear, yet the aggregation of αSyn is considered to be a prerequisite for neurodegeneration (8). It is therefore important to better understand the mechanisms of aggregation and amyloid formation to reveal new routes to ameliorate disease.

The amino acid sequence of αSyn can be divided into the N-terminal positively-charged region (residues 1-60), the highly-amyloidogenic non-amyloid-β component (NAC) (residues 61-95), and the C-terminal highly-acidic region (residues 96-140) (Fig. 1A). Sequences within all three of these regions have been revealed as essential for regulating its aggregation into amyloid (9–13). Numerous reports have also shown that truncations in the N-terminal region (14–17), can modify the propensity of αSyn to form amyloid; these changes can slow down (17) or speed up (16) amyloid formation depending on the specific truncation. In fact, single point mutations, such as Y39A or S42A in the P1 region, are sufficient to inhibit fibril formation *in vitro* and preclude aggregation in *C. elegans* (10). Post-translational modifications can also change the capacity of αSyn to form amyloid; for example, N-terminal acetylation, which occurs to αSyn in the brain (18), slows amyloid formation (19, 20). Two studies also showed that αSyn monomers bind to fibrils via their N-terminal 10-11 residues, indicating a key role of these residues in the mechanism of seeded fibril growth (21, 22). Importantly, a host of αSyn fibril structures have now been solved (23), and in the majority of cases these N-terminal 10-11 residues remain disordered and unresolved in the fibril cores.

**Figure 1.**
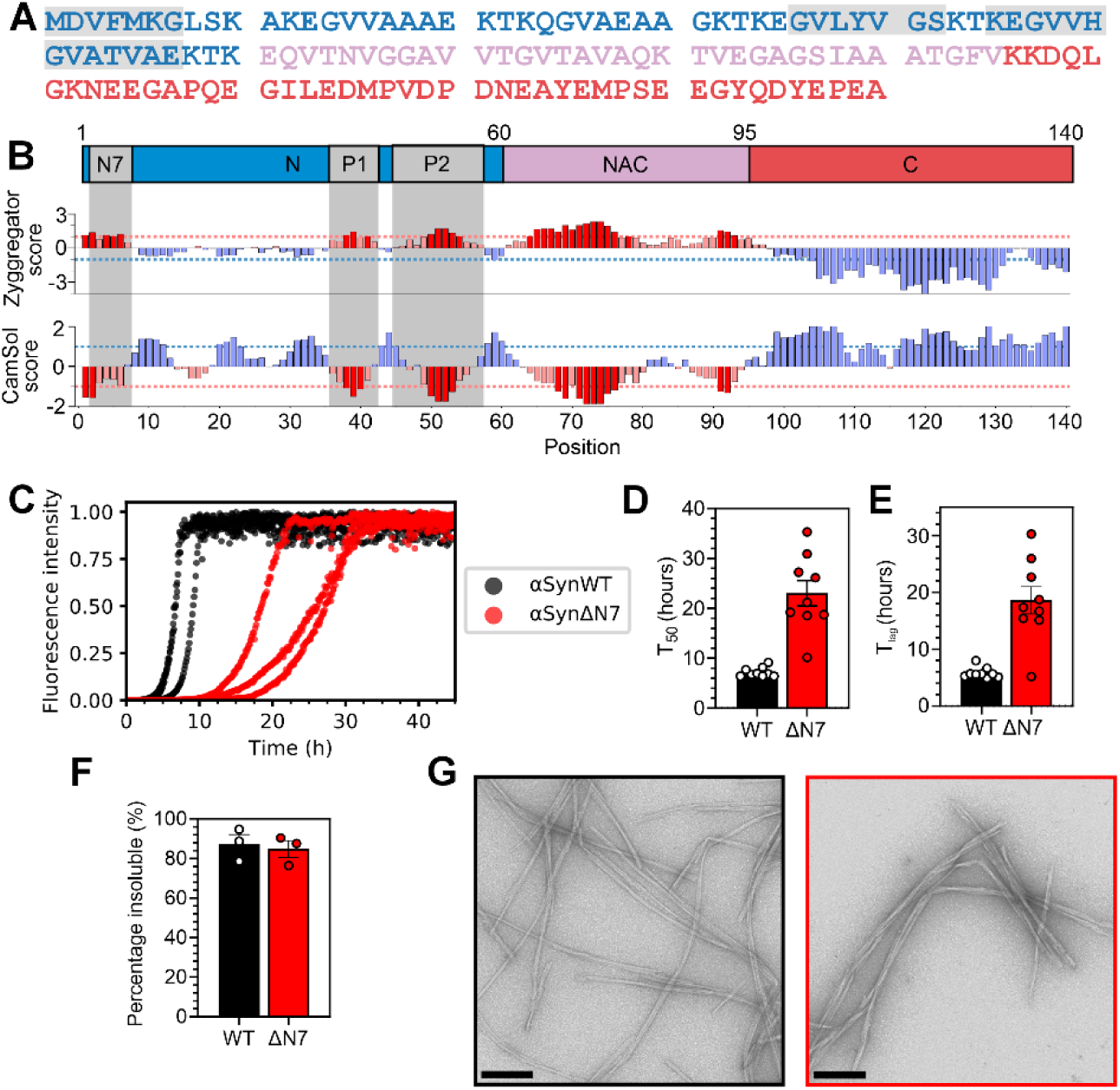
Deletion of residues 2-7 of αSyn slows amyloid formation *in vitro*. **(A)** Amino acid sequence of αSynWT. Blue = N-terminal region; pink = NAC; and red = C-terminal region. **(B)** Zyggregator (38) and CamSol (39) profiles for αSyn. Red bars indicate predicted aggregation-prone and low-solubility regions, respectively. For **(A)** and **(B)** the N7 (^2^DVFMKG^7^), P1 (^36^GVLYVGS^42^) and P2 (^45^KEGVVHGVATVAE^57^) regions are in grey. **(C)** Fibril formation kinetics of αSynWT (black) and αSynΔN7 (red). Data are normalised to maximum signal of each curve. **(D)** T_50_ and **(E)** T_lag_ values for nine replicates. **(F)** Quantification of the insoluble fraction at the endpoint of ThT assays. **(G)** Negative-stain electron micrographs of the ThT endpoints for αSynWT (black) and αSynΔN7 (red). Scale bar = 250 nm.

The N-terminal region of αSyn has also been postulated to be important for the protein’s physiological function (24–26); specifically, it is evidenced to be involved in the regulation of vesicle cycling and neurotransmitter release at presynaptic termini (24, 25). In accordance with this role, the first ∼100 residues of αSyn bind to membranes (26, 27) and alteration to the sequence of the N-terminal region can impact this interaction (9, 14). Using NMR measurements, residues 6-25 of αSyn have been proposed as the primary membrane-binding motif (28, 29), while other experiments using X-ray diffraction have shown that the N-terminal 14 residues of αSyn can insert into the membrane bilayer to form an anchor that precedes binding of additional residues (30). Furthermore, molecular dynamic simulations have suggested that the NAC domain may be involved in a double-anchor mechanism required for the physiological function of αSyn (31). Despite the differences in the proposed mechanisms of how αSyn interacts with membranes, these studies highlight the importance of the N-terminal region in mediating these interactions.

Binding of αSyn to membranes is not only important for its physiological function, but may also be critical in terms of the pathological mechanism of aggregation and Lewy body formation. Lipid membranes can trigger αSyn amyloid fibril formation (32–35), and lipids are incorporated into the fibril structures (35, 36). Together with evidence that lipids are a major component of Lewy bodies (37), this suggests that lipid-catalysed amyloid fibril formation may be involved in the aetiology of synucleinopathies.

Here, inspired by bioinformatic analyses that show residues 2-7 of αSyn to have both high aggregation propensity and low solubility (Fig. 1B) (38, 39), we show that deletion of residues 2-7 slows the rate of lipid-independent amyloid formation of αSyn *in vitro* and inhibits its recruitment to αSynWT amyloid fibril seeds. Strikingly, however, we found that deletion of these N-terminal residues does not impair the affinity of αSyn to DMPS liposomes, but abolishes lipid-stimulated fibril formation. The results are corroborated by *in vivo* findings that αSynΔN7::YFP expressed in *C. elegans* does not form significant numbers of puncta, nor does it induce the proteotoxicity associated with expression of αSynWT::YFP (40). Our findings not only develop understanding of the different regions of αSyn involved in amyloid formation, but opens the door to targeting residues 2-7 for the development of modulators of amyloid formation, potentially without affecting the functional role of the protein in membrane remodelling.

## Results

### Deletion of residues 2-7 of αSyn slows amyloid formation *in vitro*

The bioinformatic analyses conducted in our previous study that identified P1 (residues 36-42) and P2 (residues 45-57) as important regulatory regions for αSyn aggregation also identified residues 2-7 as having high aggregation propensity and low solubility (9) (Fig. 1B). Inspired by this analysis, we set out to examine the relative importance of residues 2-7 (^2^DVFMKG^7^) in αSyn amyloid formation. Accordingly, a deletion variant lacking residues 2-7 was generated, named here αSynΔN7 (note that Met1 is the initiating residue and hence was maintained in the construct). The effect of deleting these residues on the rate of amyloid formation *in vitro* (*SI Appendix*, Methods) was monitored using thioflavin T (ThT) fluorescence and the results compared with identical assays performed simultaneously using αSynWT (Fig. 1C-E). At the end of the incubation, the fraction of protein remaining in solution was determined using ultracentrifugation and SDS-PAGE (*SI Appendix,* Methods) (Fig. 1F). Additionally, negative-stain transmission electron microscopy (TEM) was used to visualise the products of the reaction (Fig. 1G). The results showed that under the conditions used (*SI Appendix*, Methods), αSynΔN7 forms amyloid-like fibrils (Fig. 1G) with similar yield (85 ± 4% and 87 ± 5% for αSynΔN7 and αSynWT, respectively) (Fig. 1F), but at a slower rate than αSynWT, with T_50_ values (time to reach 50% of the maximum ThT signal) of 23.0 ± 2.5 h and 7.2 ± 0.3 h, and lag times (T_lag_) of 18.6 ± 2.4 h and 5.8 ± 0.3 h, for αSynΔN7 and αSynWT, respectively (*SI Appendix*, Table S1). These data are consistent with a previous report on the effect of deleting residues 1-6 of αSyn on amyloid formation (17) and authenticate the predication that the N-terminal seven residues do indeed play a role in modulating the rate and/or mechanism of αSyn amyloid assembly.

The conditions used here differ to those used for our previous studies on the P1 region of αSyn (9). To compare the effects of these various deletion variants, we re-examined the amyloid formation kinetics of αSynΔP1, αSynΔP2, and αSynΔΔ (lacking both P1 and P2 regions) in the presence of a Teflon ball (*SI Appendix,* Fig. S1). The results showed that despite these more aggregation-promoting conditions, neither αSynΔP1 nor αSynΔΔ produced amyloid over the experimental timescale, while αSynΔP2 produced fibrils with a similar yield (86 ± 1.5%), but extended half-time (T_50_ = 14.8 ± 1.8 h), relative to αSynWT, as noted previously at pH 7.5 (9).

Whether deletion of residues 2-7 alters the architecture of the fibrils formed in the same buffer used for the ThT experiments was determined using cryo-EM (*SI Appendix*, Table S2). After 2D classification, a subset of twisted segments displaying a regular ∼75 nm crossover distance could be identified (Fig. S2A, B). A structure could be determined from these twisted segments to a resolution of 2.5 Å (Fig. S2C-E). The αSynΔN7 fibril core was modelled from residues 42-92 arranged in two identical protofilaments with an extensive, buried inter-protofilament interface involving residues 50-58 (Fig. S3A, B). This arrangement is consistent with those obtained previously with αSynWT and C-terminal truncations of recombinant αSyn (Fig. S3C). More extensive N-terminal deletions have been shown to lead to novel fibril architectures, for example when residues 1-40 are truncated (16). By contrast with these longer deletions, the structure determined here for αSynΔN7 suggests that while deletion of residues 2-7 slows assembly, the fibrils that result adopt a αSynWT-like fold.

Based on the similarity of cryo-EM structures of αSynWT and αSynΔN7 fibrils, we next explored the compatibility of αSynWT and αSynΔN7 in fibril co-assembly. To do this the ThT assay was repeated using each protein alone and compared with a 1:1 mixture of each monomer. The results show that when both variants are mixed the ThT kinetics closely resemble those of αSynWT alone, with the majority of the soluble protein being αSynΔN7 at the end point of the assay (Fig. S4). Hence, despite differing only by the presence of residues 2-7, αSynWT and αSynΔN7 are unable to co-assemble into amyloid.

### Residues 2-7 of αSyn are required to elongate αSynWT preformed fibril seeds

Given the surprising observation that αSynWT and αSynΔN7 do not co-assemble into amyloid when mixed, we next investigated the ability of αSynWT and αSynΔN7 fibrils to seed amyloid formation. It has been reported recently that the N-terminal 10-11 residues of αSyn monomers are required to bind to αSyn fibrils to propagate fibril growth, in a process known as seeding (21, 22). Based on this evidence, the αSynWT fibrils generated in *de novo* (unseeded) ThT assays described above (Fig. 1) were sonicated to fragment them and increase the number of fibril ends (*SI Appendix,* Methods, Fig. S5). These preformed αSynWT fibrils were then used to seed fibril elongation with αSynΔN7 monomers (10% (*v/v*) seed was added). Controls included self-seeding of αSynWT monomers with αSynWT preformed fibril seeds, self-seeding of αSynΔN7 monomers with αSynΔN7 fibril seeds, and cross-seeding αSynWT monomers with αSynΔN7 fibril seeds. Whilst αSynWT and αSynΔN7 fibrils can be rapidly elongated with the same corresponding species of monomers (self-seeding) (Fig. 2A - black and yellow, respectively, *SI Appendix*, Fig. S6, Table S3), they are distinct in their capacity to cross-seed the different monomer sequences. As expected, based on our co-assembly assay (*SI Appendix,* Fig. S4) and previous studies that mapped αSyn monomer-fibril interactions (21, 22), αSynΔN7 monomers are not recruited readily by αSynWT seeds under the experimental conditions used (Fig. 2A - blue). (Note that these experiments were performed under quiescent conditions, which do not result in fibril formation of monomers alone on the timescale employed here (Fig. 2A - green/purple)). A smaller amount of aggregated (pelletable) material resulted at the end of the experiment for αSynΔN7 monomers incubated with αSynWT fibril seeds compared with incubation of the same monomers with αSynΔN7 seeds (28 ± 10% versus 70 ± 5%, respectively) (Fig. 2B – blue and yellow, respectively) (*SI Appendix*, Table S3). In contrast, αSynWT monomers are recruited by αSynΔN7 fibrils (Fig. 2A, B – red), albeit less efficiently (i.e. fibril growth is slower than both of the self-seeding reactions (*SI Appendix*, Table S3). Notably, for this sample, fibril growth is biphasic, possibly reflecting changes in polymorphism and/or other switches of mechanism as a consequence of cross-seeding. It has been shown previously that different polymorphisms of αSyn fibrils can exhibit different intensity when bound to ThT (41) and that fibril structure can change with time (42, 43). Further work will be needed to discern the molecular mechanism underlying the biphasic kinetics observed in Fig. 2A, although we note that such behaviour cannot be explained by current two-state models of amyloid assembly (44). Together, these experiments suggest that the mechanism of recruitment and conversion to amyloid is distinct for αSynWT and αSynΔN7 monomers, with residues 2-7 being required for binding to αSynWT fibrils and conversion to a cross-β structure, but not for recruitment of αSynWT monomers to the αSynΔN7 fibril ends.

**Figure 2.**
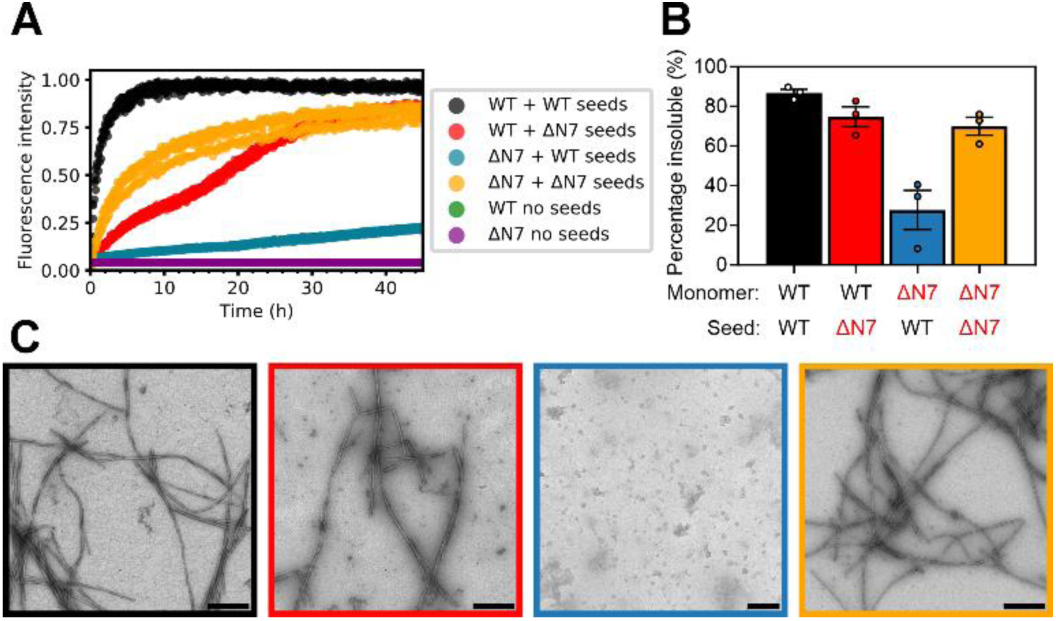
Residues 2-7 of αSyn are required to elongate αSynWT fibril seeds. **(A)** Seeded fibril growth for self- and cross-seeding of αSynWT and αSynΔN7 monomers with preformed fibril seeds (10% (*v/v*)). Data are normalised to the maximum fluorescence of the dataset. Note that for ‘WT no seeds’ (green) and ‘ΔN7 no seeds’ (purple) there is no increase in ThT fluorescence signal so the data cannot be seen readily behind each other. **(B)** Quantification of the insoluble fraction at the endpoint of ThT assays. **(C)** Negative-stain electron micrographs of the ThT endpoints, coloured as in **(B)**. Scale bar = 250 nm.

To dissect the mechanism of seeding and fibril growth further, the seeding experiments were repeated using unsonicated fibrils (*SI Appendix*, Fig. S7). The rationale for this experiment was that if elongation is the dominant mechanism of fibril assembly, the reduced number of fibril ends present in these unsonicated samples would be expected to reduce the observed rate of fibril growth. The results from this experiment showed that all reactions are slowed dramatically when fibril seeds were added without prior sonication, consistent with elongation being the dominating growth mechanism under the conditions employed (45). However, while fibrils still form in the αSynWT self-seeding reaction, the rate is substantially slower (the T_50_ is decreased from 1.2 ± 0.2 h to 14.0 ± 0.9 h with/without sonication, respectively) (*SI Appendix,* Tables S3, S4)), and no/little fibril growth is observed over the duration of the experiment for the other reaction mixtures (*SI Appendix,* Fig. S7). Importantly, this lack of fibril growth in all conditions other than αSynWT self-seeding is indicative that the presence of residues 2-7 are important in both partners (monomeric protein and fibril seeds) for successful seeding to take place. Negative-stain EM revealed that spherical oligomers result as the products of these cross-seeded reactions (both for αSynWT monomers with unsonicated αSynΔN7 seeds, and for αSynΔN7 monomers with unsonicated αSynWT seeds) (*SI Appendix,* Fig. S7D – red and blue). Similar species are observed when monomers are incubated in the absence of fibrils under these quiescent conditions (*SI Appendix,* Fig. S8). The results indicate, therefore, that residues 2-7 of αSyn are required for monomers to propagate seeded fibril growth, irrespective of whether the fibrils were created from αSynWT or αSynΔN7. Moreover, they show that failure to form fibrils (by inefficient seeding or lack of agitation in the absence of seeds) results in the accumulation of spherical oligomers that are unable to convert into a fibrillar form.

### Residues 2-7 of αSyn are not required for DMPS liposome binding

αSyn is intrinsically-disordered in aqueous solution, but has been shown to form α-helical structure upon binding to membranes (26, 27). It is widely evidenced that the N-terminal region of αSyn facilitates liposome binding (26, 28, 30), with residues 6-25 being purported to initiate binding to the lipid surface, which then induces additional residues (spanning residues 1-97) to form α-helical structure (28). Another study proposed that the N-terminal 14 residues of αSyn insert into the membrane to form an anchor (30). To determine how deletion of residues 2-7 of αSyn affects membrane binding, αSynWT and αSynΔN7 monomers were each incubated with DMPS liposomes (*SI Appendix,* Methods)) at lipid:protein molar ratios (LPRs) ranging from 0:1 to 100:1. Our rationale for focussing on synthetic DMPS liposomes, rather than a more biologically-relevant lipid mixture, is that this system is well-characterised in terms of αSyn binding affinity and ThT kinetics allowing the effects of N-terminal deletion to be compared directly with our (9) and other (34, 46, 47) previous results. Far-UV circular dichroism (CD) was also used to follow the transition of the secondary structure from unstructured to α-helical. The results showed that residues 2-7 of αSyn are not necessary for binding to DMPS liposomes (Fig. 3A, B). Binding of αSynΔN7 monomers to these membranes induces the formation of α-helical structure akin to that observed for αSynWT, resulting in a maximum of 69% and 67% helicity for αSynWT and αSynΔN7, respectively (*SI Appendix,* Methods). Fitting the resulting titration curve (Fig. 3C), as described in (34) (*SI Appendix*, Methods) yielded similar K_d_ values for lipid binding for the two proteins (1.2 ± 0.4 μM and 5.1 ± 2.3 μM for αSynWT and αSynΔN7, respectively) and a similar number of lipid molecules involved in each αSyn monomer binding event (30 ± 2 and 28 ± 5 lipid molecules per protein monomer for αSynWT and αSynΔN7, respectively). Thus, by contrast with previous reports (14), the data indicate that residues 2-7 of αSyn are not required for liposome binding, at least with the lipid type and solution conditions utilised here.

**Figure 3.**
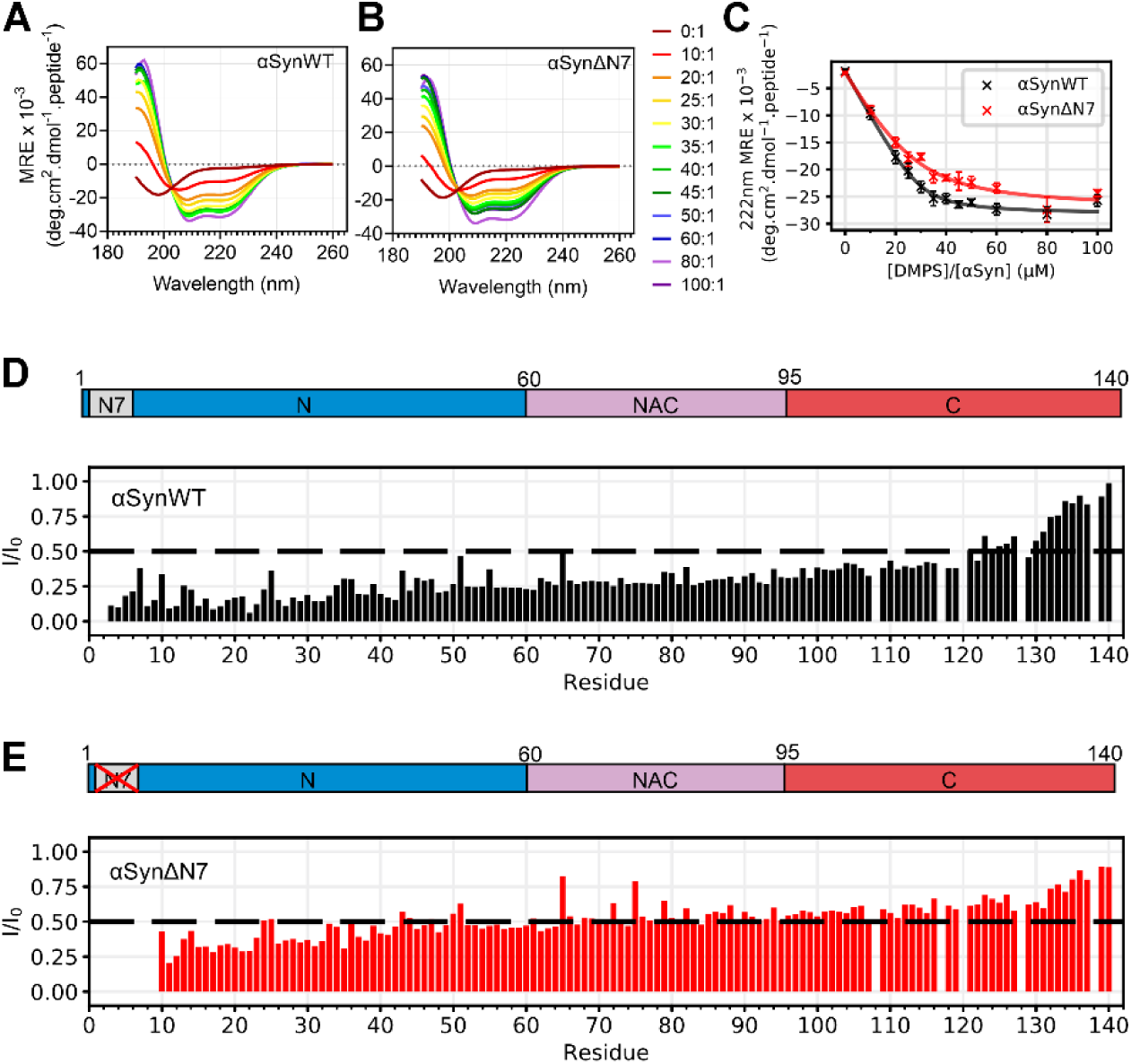
Residues 2-7 of αSyn are not necessary for DMPS liposome binding. **(A, B)** Representative far-UV CD spectra of αSynWT and αSynΔN7 as a function of the lipid-to-protein ratio (LPR) (see key). **(C)** MRE at 222 nm as a function of LPR for αSynWT and αSynΔN7. Curves were fitted using equation 6 from (34). Error bars are SEM. **(D, E)** Per-residue intensity ratios of ^1^H-^15^N HMQC NMR resonances for **(D)** αSynWT and **(E)** αSynΔN7. Spectra were collected at 30 °C at an LPR of 8:1. The dashed line indicates an I/I*_0_* of 0.5.

Given that the percentage helicity of bound αSynWT and αSynΔN7 on DMPS liposome is indistinguishable (Fig. 3C), how αSynΔN7 interacts with DMPS liposomes at a residue-specific level was next investigated using solution NMR spectroscopy (Fig. 3D, E, *SI Appendix,* Fig. S9A-C). Liposome binding was monitored by comparing the intensity of peaks in ^1^H-^15^N heteronuclear multiple quantum coherence (HMQC) spectra in the absence or presence of DMPS liposomes using an LPR of 8:1 at 30 °C (*SI Appendix*, Methods). Under these conditions, DMPS liposomes were found to be slightly above the transition temperature from gel to fluid phases (*SI Appendix*, Fig. S10). A decrease in intensity in the liposome-containing sample is indicative of binding of the protein to the membrane, with a greater loss of intensity suggesting a tighter interaction of the residue of interest with the liposomes at that site. Deletion of residues 2-7 results in higher intensity ratios for residues 10-119 for αSynΔN7 (average I/I_0_ = 0.49, standard deviation = 0.11) compared with αSynWT (average I/I_0_ = 0.28, standard deviation = 0.09) (Figure 3D, E). It is particularly remarkable that the N-terminal six residues exhibit such long-range control of membrane interaction, despite having no effect on the extent of helicity of the protein triggered by presence of the liposomes (Fig. 3A-C). To investigate if lipid ordering affects these profiles, identical experiments were performed on αSynWT and αSynΔN7 below (20 °C) and above (40 °C) the T_m_ for DMPS in the presence of αSyn (*SI Appendix,* Fig. S9D, E), (which is ∼28 °C in the presence of protein, *SI Appendix*, Fig S10). These experiments showed that the profiles differed most significantly at 30 °C, whereas below (20 °C) or above (40 °C) the T_m_ the intensity profiles for the proteins are more similar. Together this information shows that residues 2-7 of αSyn are not required for binding to these liposomes, nor do they alter liposome binding affinity, or the extent to which the αSyn molecules adopt α-helical structure, yet do affect the extent that specific residues interact with membranes under these conditions.

Overall, these results are surprising given the literature precedents that report on the importance of residues 6-25 in forming an anchor to elicit membrane binding (14, 28). The differences in results are likely a result of the specific lipid system used and/or changes in the precise regions studied, and highlight the importance of the balance of lipid type, temperature, assay method and LPR when drawing comparisons.

### Residues 2-7 of αSyn are critical for lipid-mediated fibril formation

Binding of αSyn to membranes has been shown to accelerate amyloid fibril formation by promoting heterogeneous primary nucleation (34), with lipid molecules co-aggregating with protein (33) and becoming encapsulated into the resulting fibril structures (36). Binding of αSynWT to DMPS liposomes has been shown to be sufficient to trigger amyloid formation under quiescent conditions (9, 34). To determine whether assembly of αSynΔN7 into amyloid is also stimulated by binding to membranes, the protein was incubated with DMPS liposomes and fibril formation monitored using ThT fluorescence (*SI Appendix*, Methods). The results showed that whilst a [DMPS]:[protein] ratio of 4:1, 8:1 or 16:1 results in fibril formation of αSynWT, incubation with a large lipid excess (60:1 LPR) inhibits assembly, by depleting the concentration of lipid-free monomer available for fibrillation (Fig. 4A-C), consistent with previous results (9, 34). Negative-stain EM of the αSynWT fibrils formed at an LPR of 8:1 show long, winding fibrils attached to small liposomes (Fig. 4B), as observed previously (34). Surprisingly, and in marked contrast to the behaviour of αSynWT, αSynΔN7 does not form ThT-positive amyloid fibrils at any of the lipid concentrations studied (Fig. 4D), despite binding to DMPS liposomes with similar affinity to αSynWT and forming similar helical structure in the bound state (Fig. 3A-C). Instead, spherical liposomes remain and no fibrils are observed at the end of incubation (45 h) with αSynΔN7 monomers (Fig. 4E), similar to liposomes observed after incubation in the absence of protein (*SI Appendix,* Fig. S11). Notably, when incubated at an LPR of 60:1, αSynWT and αSynΔN7 each resulted in the formation of tubulated liposomes (Fig. 4C, F), suggesting that both proteins are able to remodel the lipid bilayer by binding to the liposomes and/or integration into the lipid acyl chains. We have reported previously that an αSyn deletion variant that lacks residues 36-42 (P1 region) and 45-57 (P2 region) (named αSynΔΔ) also binds DMPS liposomes with similar affinity to αSynWT, but is unable to remodel them to form lipid tubules. Instead, smaller lipid structures are observed when incubated at an LPR of 60:1 (9). To determine whether αSynΔP1 has a similar effect on liposomes structure as αSynΔΔ, or whether deletion of the seven residue P1 segment more resembles the behaviour of αSynΔN7, the binding of αSynΔP1 to DMPS liposomes was also investigated *SI Appendix*, Fig. S12). The results were similar to those obtained for αSynΔN7, with binding resulting in helical structured (*SI Appendix*, Fig. S12A), a similar affinity (K_d_ of 4.2 ± 2.0 μM) (*SI Appendix*, Fig. S12B), and no indication of lipid-stimulated fibril formation (*SI Appendix*, Fig. S12C). Unlike αSynΔN7, αSynΔP1 forms a lower helicity in the bound state (47%, compared with the 67% helicity for αSynΔN7) (*SI Appendix*, Fig. S12A) and is unable to tubulate liposomes (*SI Appendix*, Fig. S12D). Despite their shared inability to nucleate on DMPS liposomes, the HMQC per residue intensity profiles are distinct for αSynΔP1 and αSynΔN7 at 30 °C with the former more closely resembling the profile obtained for αSynWT precluding a simple link between lipid binding and amyloid formation (*SI Appendix*, compare Fig. S12E and Fig. 3).

**Figure 4.**
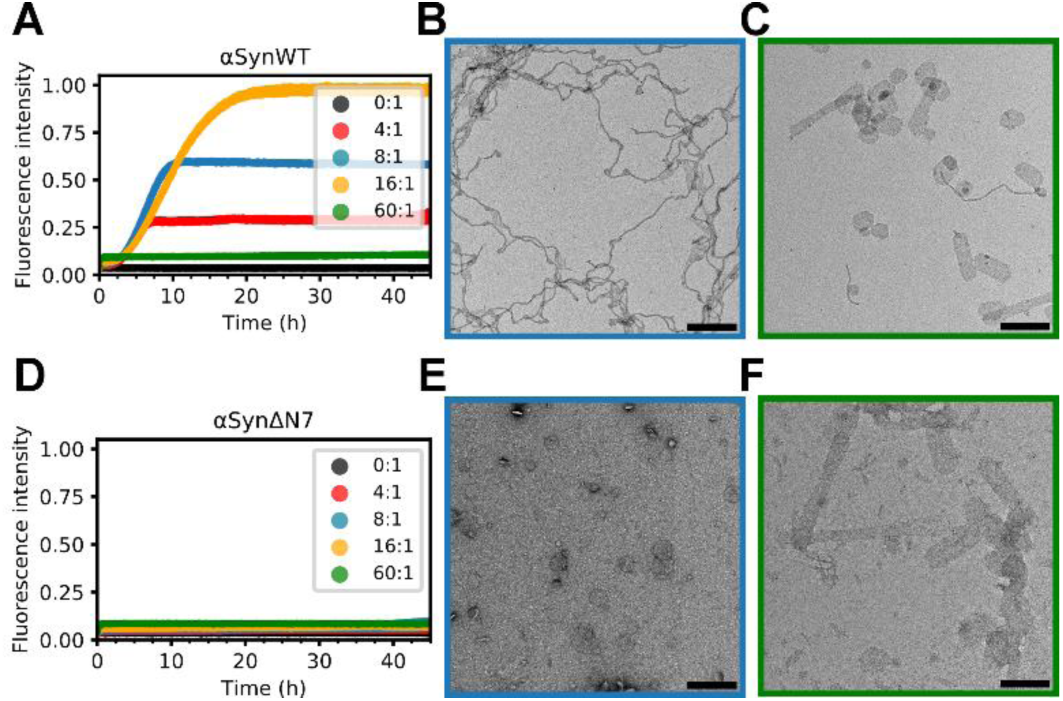
Residues 2-7 of αSyn are critical for lipid-mediated fibrillation. **(A)** Fibril formation kinetics for αSynWT in the presence of DMPS liposomes. Key indicates [DMPS]:[αSyn] ratio. Data are normalised to the maximum fluorescence intensity of the dataset. **(B)** and **(C)** Negative-stain electron micrograph of the ThT assay endpoint for [DMPS]:[αSynWT] of 8:1 and 60:1, respectively. Scale bar = 250 nm. **(D-F)**, as **(A-C)**, but for αSynΔN7.

The work above has shown that, despite binding to, and destabilising DMPS bilayers such that lipid tubulation results, αSynΔN7 lacks the ability to transform from an α-helical membrane-bound state to the cross-β structure of amyloid. This could result from the residues 2-7 being vital for this structural transition, or because these residues are required to extract the lipid from the bilayer that is a necessary step for the lipid-induced stimulation of αSynWT fibrillation (32, 33, 47).

In order to explore which residues in particular may be important for regulating lipid-induced amyloid formation we performed an alanine scan of residues 2-7 (^2^DVFMKG^7^). We used far UV CD to confirm that these mutations do not impact the α-helical propensity of αSyn in the presence of DMPS liposomes (Fig. S13), and then repeated the ThT assay with all alanine variants (Fig. S14). The results showed that whilst K6A and G7A have little effect on the rate of lipid-stimulated amyloid formation, the other variants tested (D2A, V3A, F4A, and M5A) all impede the ability of αSyn to form amyloid in the presence of DMPS liposomes. The T_50_ values of amyloid formation (*SI Appendix,* Table S5) are two-to three-fold higher for D2A (19.3 ± 0.5 h), V3A (19.6 ± 0.6 h), and F4A (16.7 ± 1.1 h), compared with αSynWT (7.2 ± 0.4 h). The single point mutation of M5A was sufficient to abolish amyloid formation on the timescale and conditions employed. This demonstrates that Met5 may be the most critical amino acid of residues 2-7 to facilitate efficient amyloid formation in the presence of these liposomes. Future work to understand the role of Met5 in lipid-induced amyloid formation will be necessary to understand the biophysical mechanisms and biological-relevance of the results presented here.

### *C. elegans* expressing αSynΔN7::YFP form fewer puncta and have no motility defects

Given that residues 2-7 of αSyn have a clear and dramatic effect on amyloid assembly *in vitro* both in the absence and presence of liposomes, the role of these residues in driving protein aggregation and its associated proteotoxicity were next tested *in vivo* using *C. elegans* as a model organism. Accordingly, a *C. elegans* strain expressing αSynΔN7::YFP in the body wall muscle cells was generated, and *in vivo* aggregation of αSynΔN7::YFP and any associated proteotoxicity (measured using motility and lifespan assays) were measured as a function of age. N2 worms, which do not express the αSyn transgene, and a strain expressing αSynWT::YFP (40) were used as controls (*SI Appendix*, Methods). Western blot analysis showed that αSynΔN7::YFP protein expression levels were similar to those of αSynWT::YFP worms, enabling their direct comparison (*SI Appendix*, Fig. S15).

Analysis of the αSynΔN7::YFP animals showed a significant reduction in the number of puncta corresponding to αSynΔN7::YFP aggregates compared with αSynWT::YFP animals at all days of adulthood measured (Day 1, 4 and 8) (Fig. 5A, B). The expression of αSynWT::YFP is proteotoxic in an age-dependent manner, with the increase of αSynWT aggregation in Day 4 adults associated with a ∼50% reduction in motility (body bends per second) compared with Day 1 adults (Fig. 5C). By contrast, expression of αSynΔN7::YFP showed no evidence of proteotoxicity in this assay throughout ageing, with motility of this strain remaining similar to N2 animals (Fig. 5C). While αSynWT::YFP expression reduced the median lifespan of the animals to 11 days of life (8 days of adulthood) compared with N2 animals which have a median lifespan of 15 days of life (12 days of adulthood), the median lifespan of αSynΔN7::YFP-expressing animals was identical to that of N2 animals (*SI Appendix*, Fig. S16, Table S6). Thus, consistent with its reduced aggregation potential observed *in vitro*, residues 2-7 are important for αSyn aggregation *in vivo* and its associated proteotoxicity.

**Figure 5.**
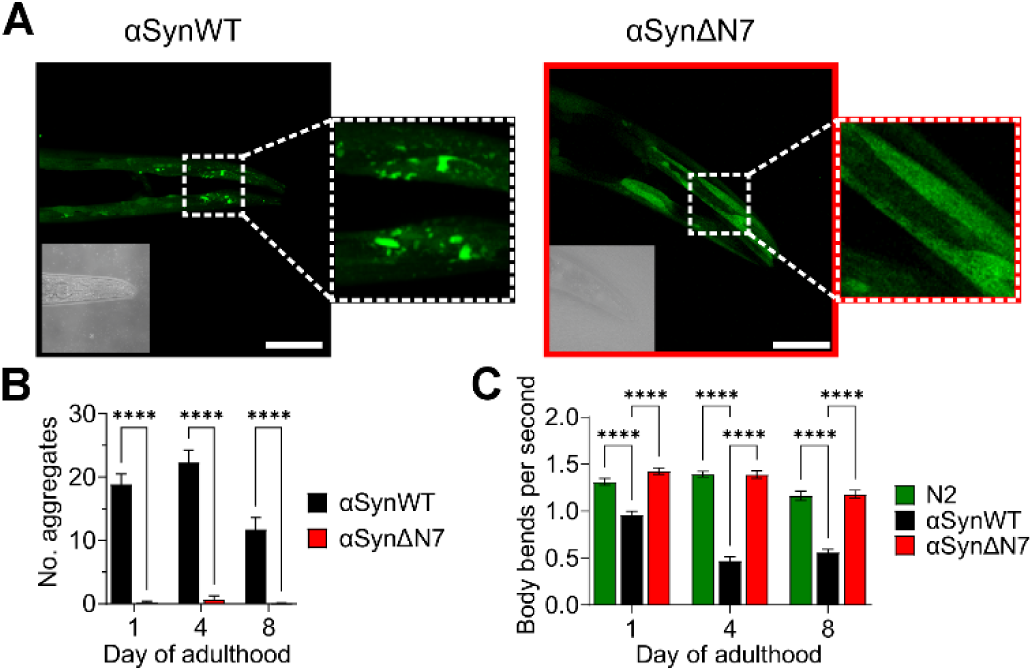
*C. elegans* expressing αSynΔN7::YFP exhibit fewer aggregates and motility defects than those expressing αSynWT::YFP. **(A)** Representative confocal images of *C. elegans* expressing αSynWT::YFP and αSynΔN7::YFP at day 8 of adulthood. Scale bar = 50 μM. Corresponding brightfield images are displayed in bottom left of each image. **(B)** Quantification of puncta in the head region of *C. elegans* expressing αSynWT::YFP or αSynΔN7::YFP. **(C)** Quantification of the motility of N2, αSynWT::YFP, and αSynΔN7::YFP animals in terms of body bends per second. (****) = p < 0.0001.

## Discussion

### Residues 2-7 of αSyn control amyloid formation *in vitro*

The N-terminal region of αSyn is known to be important for its interaction with membranes, and to play a role in determining the ability of the protein to assemble into amyloid fibrils, both in the presence (9, 48) and absence of a membrane (16, 17). However, there is conflicting evidence as to the precise role of this region in assembly (16, 17). Previous studies have shown that truncation of the N-terminus by 13, 35 or 40 residues accelerates amyloid assembly, and results in the formation of fibrils with a distinct structure to those observed for αSynWT (16). The N-terminal region has also been shown to be important for binding of monomers to the fibril surface to propagate seeded fibril growth, with truncation of 40 or more residues from the N-terminus disabling the ability of monomers to be recruited by αSynWT fibrils, while smaller truncations have no effect (16).

Here we have focussed on the role of a short, six amino acid region (residues 2-7) of αSyn on amyloid assembly, inspired by bioinformatics analyses which suggest that this region has both high aggregation potential (Zyggregator (38)) and low solubility (CamSol (39)) (Fig. 1A, B). Despite these features, residues 2-7 do not form part of the ordered fibril core, but remain dynamically disordered (23). Interestingly the N7 region has similar properties judged by these algorithms to a second region in the N-terminal domain (residues 36-42 (named P1)), which we previously showed to be essential for *de novo* fibril formation at neutral pH *in vitro* (9, 10) and is involved in forming the core of most αSyn fibril structures (23).

Via a systematic study of *de novo* (unseeded) and seeded assays we show here that deletion of residues 2-7 of αSyn has a very different effect on fibril growth compared with the large N-terminal truncations described above, with deletion of these six residues slowing amyloid formation *de novo* (T_lag_ and T_50_ are increased ∼3-fold) (Fig. 1D, E) (*SI Appendix,* Table S1). We were able to resolve a 2.5 Å resolution cryo-EM structure (*SI Appendix,* Fig. S3) which shows a typical αSynWT fibril fold and architecture, suggesting that deletion of residues 2-7 does not significantly alter the final fibrillar state. This highlights the critical sensitivity of fibril growth on the precise sequence of the N-terminal region, with acceleration or retardation of fibril assembly being dependent on the location of the amino acid deletions or substitutions made, as well as the solution conditions (9, 10, 16, 17). Such sensitivity also rationalises why truncations in the N-terminal region of αSyn are found in Lewy bodies in Parkinson’s patients (49), why post-translational modifications in the N-terminal region (such as N-terminal acetylation (19, 20)), and mutations associated with familial disease (many of which lie in the N-terminal region) (2–6, 50–54) often (A30P, E46K and A53T/V/E), but not always (H50Q and G51D), enhance the rate of amyloid formation (48). Whilst we focus here on its non-N-terminally acetylated version, αSyn is natively N-terminally acetylated (18). How this post-translational modification affects amyloid formation of αSynΔN7 will require further work. Changes in the C-terminal region can also modulate amyloid formation, with truncations in this region (e.g. 1-103 or 1-119) (12), metal ion binding (55), and post-translational modifications (56, 57) having profound, and often very different effects, on amyloid formation, at least *in vitro*.

### Lipid-stimulated amyloid formation

The second unexpected observation we report is that the affinity of αSynΔN7 monomers for DMPS liposomes is not affected by deletion of residues 2-7, by contrast with studies which suggest that this region is essential for lipid binding and membrane insertion (14, 29, 30). This highlights an extreme sensitivity of the properties and effects of lipid interaction dependent on αSyn sequence, lipid type and reaction conditions. We also show that the extent of α-helix formation upon lipid binding is not affected by deletion of residues 2-7 (Fig. 3A-C), yet this deletion does impede the extent to which residues interact with the liposomes (Fig. 3D, E).

We have shown that amyloid formation of αSynΔN7 is not stimulated by liposome binding (Fig. 4), in marked contrast to αSynWT, wherein lipid acts as a substrate for its assembly into amyloid, with the fibril yield and rate of assembly depending on the LPR (33–35). Lipids can become integrated into the core of the cross-β amyloid fold (33), being clearly observed in the cryo-EM structures of the resulting fibrils (36). A model by which αSyn extracts lipid from membranes to enable αSyn-lipid co-aggregation to occur has been proposed (32).

We postulated based on these observations, that αSynWT and αSynΔN7 may have different effects on the stability of lipid molecules within the DMPS bilayer. However, and again surprisingly, using DPH anisotropy, we showed that this is not the case (*SI Appendix,* Fig. S10), with both proteins affecting the bilayer to a similar extent, and both monomers remodelling the liposomes resulting in tubulation (Fig. 4C, F). One possible explanation of the different outcomes of lipid binding for the two proteins, could be that lipid extraction, which is a prerequisite for αSyn-lipid co-aggregation and amyloid fibril formation, can no longer occur efficiently upon deletion of residues 2-7. The results from our alanine scan show that the most important residue of the N7 region to elicit lipid-induced amyloid formation is Met5, with other residues playing a minor (Asp2, Val3, and Phe4) or no role (Lys6, Gly7) (Fig. S14). We hypothesise that these residues – particularly Met5 – are important for utilising the lipid as a substrate, facilitating the conversion from the lipid-bound α-helical protein to the cross-β structure of amyloid. Further experiments will be needed to test this hypothesis and to better understand how and why lipid-induced fibrillation of αSynΔN7 and αSynM5A does not result upon lipid binding.

Perhaps the most striking, and biologically relevant, result from the investigations described above is the difference in aggregation-propensity of αSynWT::YFP and αSynΔN7::YFP in the *C. elegans* model organism. The strain expressing αSynΔN7::YFP forms few puncta and exhibits none of the proteotoxic effects that are observed in strains expressing αSynWT::YFP. Indeed, animals expressing αSynΔN7::YFP were indistinguishable from N2 control worms (which do not express any form of αSyn) in terms of motility and lifespan.

### Summary and outlook: Function versus aggregation

Given that residues 2-7 are highly conserved in the synuclein family (the sequence is identical in αSyn and βSyn, and has a single amino acid substitution (^2^DVFKKG^7^) in γSyn), the question arises as to why these residues persist through evolution and in different synuclein family members. Given the difference in residue 5 between αSyn (Met) and γSyn (Lys), it is particularly interesting that it was recently shown that γSyn is unable to undergo lipid-induced amyloid formation within 35 hours of the experiment (58). This supports the findings discussed above, that Met5 is an important residue in the mechanism of lipid-induced amyloid formation. The physiological function of αSyn is widely evidenced to involve its N-terminal domain binding to membranes at the pre-synaptic terminals and remodelling these membranes to promote synaptic vesicle docking (24) and neurotransmitter release (25).

We have shown that under the conditions tested here, deletion of residues 2-7 of αSyn does not perturb the affinity of the protein for DMPS membranes (Fig. 3A-C). Nor does loss of these residues affect the protein’s ability to reduce lipid T_m_ (*SI Appendix*, Fig. S10), or to remodel liposomes, causing their tubulation (Fig. 4C, F). Nonetheless, deletion of residues 2-7 does affect the residues involved in liposome binding (Fig. 3D). Together, these results suggest that while residues 2-7 enhance the amyloidogenic potential of αSynWT, these residues may not be essential for the physiological function of αSyn *in vivo* (although further work using a more biologically-relevant lipid system is needed to substantiate this claim). The region 2-7 of αSyn thus provides a new and exciting opportunity to develop inhibitors of amyloid assembly that target the N-terminal region, yet may not affect the functional role of the protein in membrane remodelling at the synapse. It is hoped that the results presented not only here, along with other αSyn mutational studies (59,60), when combined via the powers of artificial intelligence tools, may enable better understanding the contributions of different residues and regions of the protein sequence in its aggregation and function, such that critical targets can be found and inhibitors developed for the prevention and treatment of disease.

## Materials and Methods

Detailed explanations regarding recombinant protein expression and purification, ThT assay conditions, quantification of insoluble material, TEM, cryo-EM, liposome generation, CD, NMR, measurements of lipid dynamics, electron microscopy, and *C. elegans* strain generation, imaging and behavioural experiments can be found in the *SI Appendix*.

## Data Availability

The data associated with this paper are openly available from the University of Leeds Data Repository. https://doi.org/10.5518/1422 (61). The cryo-EM map and model for αSynΔN7 are deposited to the PDB and EMDB respectively with codes 8QPZ and 18570.

## Acknowledgments

We thank members of the Radford and Brockwell labs for insightful discussions and support. We thank Alexander Taylor, Emily Byrd, and Leon Willis for many helpful contributions and discussions, and Tabitha Howe for technical support. Negative-stain EM, CD and NMR data were collected using facilities in the Astbury Biostructure Laboratory and were funded by the University of Leeds and the Wellcome Trust (094232/Z/10/Z). Confocal microscopy data were collected with instrumentation from the Leeds Bioimaging Facility (Wellcome Trust funded (104918MA)). K.M.D. and M.W are funded by MRC (MR/N013840/1 and MR/T011149/1). B.R. and J.A.C. by BBSRC (BB/W007649/1 and BB/T008059/1), J.M.M., S.M.U. and D.T. by Wellcome (222373/Z/21/Z, 215062/Z/18/Z, and 204963), S.E.R. holds a Royal Society Professorial Research Fellowship (RSRP/R1/211057). For the purpose of open access, the authors have applied a CC BY public copyright licence to any Author Accepted Manuscript version arising from this submission.

## Author Contributions

K.M.D. prepared *in vitro* samples and measured ThT fluorescence data. K.M.D. performed all experiments on liposomes. K.M.D. performed all *C. elegans* experiments. B.R. carried out and analysed NMR experiments. J.A.C and J.M.M. carried out lipid fluidity measurements. D.T. and M.W. performed cryo-EM experiments. S.M.U. provided preliminary ThT fluorescence data. G.N.K assisted with CD experiments. N.A.R. advised on cryo-EM experiments, P. vO.H. advised on experiments involving *C. elegans*. S.E.R. and D.J.B. developed the ideas and supervised the work. All authors contributed to the preparation of the manuscript.

