## Supplementary Information for "Residues 2-7 of α-synuclein regulate amyloid formation via lipid-dependent and -independent pathways"

**This PDF file includes:**

Supporting text, Methods

Figures S1 to S16

Tables S1 to S6

### SI Appendix, Materials and Methods

#### αSynΔN7 and alanine scan mutants molecular cloning

The plasmids for αSynΔN7 and alanine scan mutants used for recombinant protein expression in *E. coli* was derived from a pET23a vector encoding αSynWT (kindly provided by Professor Jean Baum, Department of Chemistry and Chemical Biology, Rutgers University, NJ, USA). Q5 site-directed mutagenesis (NEB), followed by a kinase, ligase, DpnI (KLD) treatment was carried out on the αSynWT plasmid. Primers to generate αSynΔN7 were designed to delete the DNA sequence corresponding to residues 2-7 ($^{2}$DVFMKG$^{7}$) of αSyn. Primers for the alanine scan mutants were designed such that the amino acid at each of the positions from 2-7 would be mutated to an alanine. Successful deletion/mutation was confirmed by transformation of *E. coli* DH5α cells, Miniprep (Qiagen), and subsequent sequencing (Source Bioscience).

#### Recombinant protein expression and purification

Competent *E. coli* BL21 (DE3) cells were transformed by heat-shock at 42 °C with the pET23a vectors discussed above. Cells were plated on 100 µg/mL carbenicillin LB agar plates and grown at 37°C overnight. A 100 mL LB medium starter culture containing 100 µg/mL carbenicillin was inoculated with a single colony and incubated at 37 °C, 200 rpm shaking overnight. For $^{15}$N-labelled protein, 100 mL LB was inoculated (with 100 µg/mL carbenicillin) and eight hours later cells were pelleted and resuspended in 500 mL HCDM1 medium (10 g $K_2HPO_4$, 10 g $KH_2PO_4$, 7.5 g $Na_2HPO_4$, 9 g $K_2SO_4$, 1 g $^{15}$N-labelled $NH_4Cl$ per 1 L culture, supplemented with 2mM $MgCl_2$, 100 µM $CaCl_2$, 4 g/L glucose and 100 µg/mL carbenicillin) such that the starting $OD_{600}$ = ~ 0.04. The culture was then incubated at 37 °C, 200 rpm shaking overnight. The following day, 1 L medium (LB or HCDM1) was inoculated with 15 mL starter culture and grown at 37 °C, 200 rpm shaking. When the culture reached $OD_{600}$ of 0.6, protein expression was induced with addition of 1 mM isopropyl-β-D-thio-galactopyranoside (IPTG). Cultures were then grown for an additional 4 h before harvesting the cells.

Cells were harvested at 5,000 r.p.m. for 30 min (rotor JA 8.1.) at 4 °C. The cell pellet was then resuspended in lysis buffer (20 mM Tris-HCl pH 8.0, 2 mM $MgCl_2$, 5 mM DTT, 1 mM PMSF, 2 mM benzamidine, 100 µg/ml lysozyme, and 20 µg/ml DNase) using 25 mL lysis buffer per 1 L culture equivalent. The cell pellet was homogenised and incubated for 30 min on a roller at room temperature. The lysates were then heated at 80 °C for 10 min before centrifugation at 35,000 x *g* for 30 min. 30 % (*w/v*) ammonium sulphate was added to the resulting supernatant fraction and left to incubate on a roller for 30 min at 4 °C to precipitate the protein. The protein was then pelleted at 35,000 x *g* for 30 min at 4 °C and the supernatant was discarded. The 30 % (*w/v*) ammonium sulphate precipitation step (30 min, 4 °C) and subsequent centrifugation step (35,000 x *g*, 30 min, 4 °C) was repeated and the remaining pellet was stored at -20 °C until further processing.

The protein pellet was resuspended in 20 mM Tris-HCl, pH 8.0 wash buffer (1 L culture equivalent was resuspended in 100 mL buffer). The resuspended protein was loaded onto a manually packed Q-Sepharose anion exchange column (~80 mL). The column was washed with two column volumes (CV) of wash buffer, before eluting the protein using a gradient of 0-500 mM NaCl over two CV. The column was then washed with two CV 500 mM NaCl then two CV 1 M NaCl. The αSyn-containing fractions were pooled and dialysed into 5 mM ammonium bicarbonate. The partially-purified protein was lyophilised and stored at -20 °C until further processing.

Next, the protein was resuspended in PBS buffer (137 mM NaCl, 2.7 mM KCl, 8.1 mM $Na_2HPO_4$ and 1.5 mM $KH_2PO_4$; pH 7.4) at a concentration of 2 mg/mL and loaded onto a HighLoad™26/60 Superdex 75 pg gel filtration column. 5 mL injections were loaded using a 50 mL superloop. Peaks containing αSyn were pooled and dialysed into 5 mM ammonium bicarbonate, then lyophilised and stored at -20 °C.

**Thioflavin T (ThT) assay**

Lyophilised protein was dissolved in PBS at an approximate concentration of 10 mg/mL. Resuspended protein was centrifuged at 16,000 r.p.m. for 30 min at 4 °C to remove insoluble material. The protein concentration was then determined using the $A_{280}$ and $\varepsilon$ = 5,960 $M^{-1}$ $cm^{-1}$ for both αSynWT and αSynΔN7. To measure *de novo* fibrillation, αSyn protein (100 μM) and ThT (20 μM) were mixed and 100 μL was added per well of a 96-well non-binding flat-bottom assay plate (Corning). A 3 mm-diameter Teflon ball (PolySciences) was added to each well and the ThT assay was carried out at 37 °C, 600 rpm orbital shaking in FLUOstar Omega Plate Reader (BMG Labtech). ThT fluorescence was then monitored using excitation at 444 nm and fluorescence emission was monitored at 480 nm. Each experiment was repeated a minimum of three times, with three replicates per experiment.

For ThT experiments with sonicated seeds, fibrils taken from the endpoint of the *de novo* fibril growth experiments were sonicated on ice using a Cole-Parmer-Ultraprocessoer-Sonicator twice for 30 sec at 40% maximum power with a 30 sec break in between. Sonicated or unsonicated fibril seeds were then added to monomeric protein (50 µM) at a seed concentration of 10% (*v/v*). Fibril growth was allowed to proceed for a further 45 h 37 °C under quiescent conditions in FLUOstar OPTIMA Plate Reader (BMG Labtech).

Lag times ($T_{lag}$) were calculated by first fitting the following equation from (1):

$$Y = y_i + m_i t + \frac{y_f + m_f t}{1 + e^{-[(t-t_{50})/\tau]}}$$

where $Y$ is the fluorescence intensity over time ($t$). $y_i$ and $y_f$ are the y-intercepts of the initial and final baselines. $m_i$ and $m_f$ are the slopes of the initial and final baselines, respectively. $t_{50}$ is the time taken to reach 50% of the elongation phase and $\tau$ is the elongation time constant. The $T_{lag}$ was then calculated using:

$$T_{lag} = t_{50} - 2\tau$$

Error bars on $T_{50}$ and $T_{lag}$ plots are standard error of the mean. To measure liposome-mediated fibrillation kinetics, αSyn (50 µM) was incubated with DMPS liposomes (at molar ratios indicated in the text) in 20 mM sodium phosphate buffer (pH 6.5). 100 µL was added per well of a 96-well non-binding flat-bottom assay plate (Corning). The plate was incubated at 30 °C in FLUOstar OPTIMA Plate Reader (BMG Labtech) under quiescent conditions.

**Pelleting assay**

To quantify the amount of soluble protein remaining after each fibril growth assay, the samples were centrifuged at 100,000 x *g* for 30 min at 4 °C to separate soluble and insoluble fractions. Whole and soluble fractions were then diluted to 8.33 µM and 15 µL loaded onto a 15% Tris-Tricine SDS-PAGE gel. The gel was stained with InstantBlue Coomassie stain and imaged on the Alliance Q9 Imager (Uvitec). Densitometry was carried out using the Nine-Alliance software. Error bars on figures are standard error of the mean.

**Negative stain transmission electron microscopy (TEM)**

At the end of each ThT assay, samples were applied onto an in-house prepared carbon-coated copper grid. The grid was then washed three times with 18 MΩ $H_2O$ and negatively stained with 2 % (*w/v*) uranyl acetate twice. Images were collected on an FEI Tecnai T12 electron microscope.

### Cryo-EM data collection

100 µM of αSynΔN7 in PBS was incubated in a thriller shaker for a week at 37 °C, 600 rpm for fibrillation. 4 µL was then applied onto 60s plasma cleaned (GloCube, Quorum) Lacey carbon 300 mesh grids. The sample was blotted and frozen in liquid ethane using a Vitrobot Mark IV (FEI) with a 0.5 s wait and 5 s blot time respectively. The Vitrobot chamber was maintained at close to 100% humidity and 4 °C. The cryo-EM dataset was collected at the University of Leeds Astbury centre using a Titan Krios electron microscope (Thermo Fisher) operated at 300 kV with a Falcon IV detector (Thermo Fisher) and Selectris energy filter set with a 10 e-V slit width (Thermo Fisher). A nominal magnification of 130,000x was set yielding a pixel size of 0.95 Å. A total of 5,464 movies were collected with a nominal defocus range of -1.4 to -2.6 µm using a total dose of ~45 e-/Å2 over an exposure of 4.59 s, which corresponded to a dose rate of ~8 e-/pixel/s. Each movie was collected as EER frames.

### Cryo-EM data processing

Each movie stack was aligned, dose weighted and summed using motion correction in RELION4 (2) and CTF parameters were estimated for each micrograph using CTFFIND v4.14 (3). Fibrils from around a hundred micrographs were manually picked in RELION and the extracted segments used to train automated filament segment picking with Topaz (4). A total of 315,777 helical segments were extracted 3x binned in RELION (box dimensions of ~76 nm) for the initial rounds of 2D classification, after which the selected segments were re-extracted un-binned (box dimensions of ~47 nm) for the third round of 2D classification. Throughout, the VDAM classification algorithm was used to separate out picking artefacts to leave 163,259 fibril segments for further processing. The αSynΔN7 dataset was split into fibrils with (100,454) and without (62,805) distinct crossovers.

For starting 3D classification, an initial 3D template was generated from a single unbinned 2D class average using the relion_helix_inimodel2d command (5) along with measured helical crossover estimates from 3x binned 2D class averages. The first 3D classification run with the initial model used a fixed twist (based on the updated estimate) and rise (4.80 Å), with a t-value of 30, 1.8˚ sampling, and a strict high-resolution limit of 4 Å with 3 output classes. For the second round of 3D classification, helical searches of the twist were employed with a fixed rise (4.80 Å) with a t-value of 30, 0.9˚ sampling and a strict high-resolution limit of 4 Å with 3 output classes. In the third 3D classification run, narrow helical searches of both the twist and rise were employed with a t-value of 30, 0.9˚ sampling and a strict high-resolution limit of 4 Å with two output classes. The most ordered class was selected from this final classification to move on to refinement.

A couple of sequential rounds of 3D refinement were needed to get the halfmaps to converge before CTF refinement of the per-particle defocus estimates, Bayesian polishing and then the final run of 3D refinement. Narrow helical searches of twist and rise were used up until the final refinement, with t-values of 15, initial sampling of 0.9˚ and initial lowpass-filtering of 6 Å. Post-processing with a soft mask (extended by 4 pixels and soft-edge for 6 pixels, z length of 20%) was used to obtain gold-standard (at an FSC value of 0.143) resolution estimate of the final map. The final αSynΔN7 map is at a resolution of 2.5 Å. The full collection and processing details are shown in *SI Appendix,* Table S2.

**Model building and refinement**

After docking several different published recombinant αSyn fibril structures, PDB: 6osl (6) was selected as a starting model for building in Coot (7). One chain of the template was rigid-body fitted into the density and local regions adjusted where they deviated from the map. Both Ramachandran and rotamer outliers were monitored and minimized during building in Coot. The corrected chain was copied and rigid body fit to generate a 12-chain model for 6 layers of the fibril core followed by real space refinement against the postprocessed map in Phenix v1.17.1 (8). NCS restraints were applied to prevent divergence of repeating chains in the layers along the fibril axis. The final refined model was assessed using MolProbity (9) and deposited, with the final model statistics summarised in *SI Appendix,* Table S2.

**Reversed-phase High Performance Liquid Chromatography (RP-HPLC)**

RP-HPLC was used to resolve αSynWT and αSynΔN7 from the soluble fraction of a co-aggregation ThT assay using a Shimadzu UK Ltd. Nexera LC-40 system. The Nucleosil 300 C4 column (Chromex Scientific Ltd.) was equilibrated with 95 % solvent A (0.1 % (*v/v*) trifluoroacetic acid (TFA)) and 5 % solvent B (acetonitrile and 0.1 % (*v/v*) TFA) at a flow rate of 1 mL/min. The sample of the endpoint from the co-incubated ThT assay was centrifuged at 100,000 x *g* for 30 min at 4 °C and 30 μL of the soluble fraction was injected onto the column from a polypropylene HPLC vial (Thermo Fisher). For the monomeric αSynWT and αSynΔN7 controls, lyophilised protein was freshly dissolved in PBS and resuspended protein was centrifuged at 16,000 r.p.m. for 30 min at 4 °C to remove insoluble material. Samples were eluted over an increasing gradient of solvent B (5 – 80 %) over 10 mins, then 95 %

solvent A for 5 mins. Protein was detected at a wavelength of 280 nm by photodiode array detection. The peaks were identified using LabSolutions software provided with the instrument.

**Generation of liposomes**

DMPS lipids (Avanti) stored at -20 °C were removed from the freezer and allowed to reach room temperature. Lipids were dissolved in 20% (*v/v*) methanol and 80% (*v/v*) chloroform. One or two days before each experiment, the lipids were desiccated using a 45 °C water bath under a gentle stream of $N_2$ gas, and placed in a vacuum chamber overnight. The following day, desiccated lipids were resuspended in 20 mM sodium phosphate buffer, pH 6.5 at a concentration of 40 mM. Lipids were extruded using a 100 nm membrane. Dynamic light scattering (DLS) was used to calculate the radius of the liposomes to be (~64 ± 1 nm).

**Circular dichroism (CD)**

CD experiments were carried out using an Applied Photophysics Chiroscan spectrometer. Spectra were acquired using a 2.0 nm bandwidth and 1.0 mm path length cuvettes. All CD spectra were acquired at 30 °C. αSyn monomer was dissolved in 20 mM sodium phosphate buffer (pH 6.5) at a concentration of 25 μM and incubated with liposomes (at LPRs ranging from 0:1 to 100:1) for 3 min at 30 °C. Spectra for each sample were acquired three times, and data were blank-corrected, averaged and converted from mdeg to mean residue ellipticity (MRE) values. This experiment was repeated three times.

Binding affinity ($K_d$) and stoichiometry values (L) were calculated by fitting the equation described in (10):

$$X_B = \frac{\left([\alpha Syn] + \frac{[DMPS]}{L} + K_d\right) - \sqrt{\left([\alpha Syn] + \frac{[DMPS]}{L} + K_d\right)^2 - \frac{4[DMPS][\alpha Syn]}{L}}}{2[\alpha Syn]}$$

where $X_B$ is the fraction of αSyn bound to the membrane and can be expressed as:

$$X_B = \frac{CD_{obs} - CD_F}{CD_B - CD_F}$$

where $CD_{obs}$ is the observed CD signal at 222 nm, and $CD_F$ and $CD_B$ are the CD signals of free and bound αSyn, respectively.

To calculate the percentage helicity of αSynWT and αSynΔN7, the following equation from (11) was used:

$$Percentage\ helicity = 100 \times \frac{\theta_{222nm}^{exp} - \theta_{222nm}^{u}}{\theta_{222nm}^{h} - \theta_{222nm}^{u}}$$

where $\theta_{222nm}^{u}$ and $\theta_{222nm}^{h}$ correspond to the ellipticity of a protein with 0% and 100% helical content, respectively. These values have been estimated to be −3,000 deg.cm$^2$.dmole$^{-1}$ ($\theta_{222nm}^{u}$) and −39,500 deg.cm$^2$.dmole$^{-1}$ ($\theta_{222nm}^{h}$). $\theta_{222nm}^{exp}$ is the observed ellipticity under saturating conditions.

**NMR experiments**

2D ($^1$H, $^{15}$N) heteronuclear multiple quantum coherence (HMQC) spectra were recorded at 20 °C, 30 °C and 40 °C using a AVANCE III Bruker spectrometer (600 MHz) equipped with a triple channel QCI-P cryoprobe. Spectra were recorded on samples of 25 µM [U-$^{15}$N] WT αSynWT, αSynΔN7 or αSynΔP1 in 20 mM sodium phosphate buffer, pH 6.5, 10% (*v/v*) $D_2O$ in the absence or presence of DMPS liposomes (~100 nm diameter) at an LPR of 8:1. All spectra were processed using Topspin 3.7.0 and analysed using CCPN analysis software (12). Published assignments of αSynWT (BMRB 16543) (13) were transferred to the data for αSynΔN7 and αSynΔP1 at 20 °C. Assignments could then be transferred to spectra collected at higher temperatures by following peak positions at different temperatures and assuming a linear relationship between temperature and chemical shift for a given peak (14). Peak heights were used to calculate intensity ratios for peaks in the absence and presence of lipid.

**DPH assay**

1,6-diphenyl-1,3,5-hexatriene (DPH) was dissolved at room temperature by stirring overnight in 100% ethanol to make a stock concentration of 2 mM. DPH was then added to DMPS liposomes at a molar ratio of 1:600 [DPH]:[DMPS] and the mix was placed at 45 °C for 30 min to allow incorporation of the DPH into the lipid bilayer. This dye integrates into the lipid acyl chains and reports on lipid fluidity via measurement of its fluorescence polarisation (15). The DMPS-DPH mix was then added to αSynWT or αSynΔN7 monomers (final concentration of 50 µM protein), and a final LPR of 60:1 or 8:1. The final concentration of ethanol in the sample was 0.025% (*v/v*). Fluorescence polarisation was measured on a PTI Quantamaster Series fluorometer. DPH fluorescence was excited at 355 nm with fluorescence emission collected at 430 nm. The fluorescence polarisation was acquired at 1 °C intervals from 10 to 60 °C, with a 2 min equilibration at each temperature and a 5 sec integration time. Milli polarisation (mP) was calculated by:

$$mP = 1000 \times \frac{I_{VV} - G \times I_{VH}}{I_{VV} + G \times I_{VH}}$$

where $I_{VV}$ is the measured intensity of emitted fluorescence with the excitation polarisers in the vertical (i.e. 0 °) position and the emission polariser also in the vertical position. $I_{VH}$ is the same as $I_{VV}$, but with the emission polariser in the horizontal position (i.e. 90 °). G is the G factor, given as:

$$G = \frac{I_{HV}}{I_{HH}}$$

where $I_{HV}$ is the measured fluorescence intensity with the excitation polariser in the horizontal position and emission polariser in the vertical position. $I_{HH}$ is the same as $I_{HV}$ but with the emission polariser in the horizontal position.

**_C. elegans_ strain generation and maintenance**

The NL5901 strain (expressing *unc-54p::αSynWT::*YFP) and the pPD30.38 vector encoding *unc-54p::αSynWT::YFP* was kindly provided by Professor Ellen Nollen (University of Groningen, Netherlands). The *unc-54p::αSynΔN7::YFP* strain was generated by Gateway® cloning (Invitrogen). The BP reaction was used to generate the *αSynΔN7* entry clone. For the LR reaction, the entry clones containing *αSynΔN7* was combined with pDONR™P4-P1R and pDONR™ P2RP3 vectors containing *unc-54p* and *YFP*, respectively. The product of the LR reaction was used to transform DH5α cells and the expression vector was checked by sequencing. This construct was used to generate the transgenic *C. elegans* strains by microinjection into the N2 nematodes, resulting in the generation of the strain PVH247. *C. elegans* strains were maintained using standard methods at 20 °C with *E. coli* OP50-1 as a food source.

**Western blotting to quantify protein expression levels**

~400 *C. elegans* worms were collected and placed into M9 solution. Nematodes were pelleted by a brief centrifugation, and washed with M9 solution three times. The remaining pellet was resuspended in 20 µL lysis buffer (20 mM Tris-HCl, pH 7.5; 10 mM β-mercaptoethanol; 0.5% (*v/v*) Triton X-100; supplemented with complete protease inhibitor (Roche)). The samples were snap-frozen in liquid nitrogen and freeze-thawed three times. The *C. elegans* worms in the samples were then lysed on ice using a motorised pestle. The samples were then centrifuged at 1000 x *g* for 1 min and the supernatant containing the worm lysate was collected. The protein concentration of each sample was calculated using a Bradford assay (Thermo Fisher). Lysates were diluted to 1 μg/μL and mixed with 2x SDS loading buffer (2 % (*w/v*) SDS, 10 % (*v/v*) glycerol, 0.1 % (*w/v*) bromophenol blue, 100 mM DTT) and boiled for 10 min. 15 μg of protein was loaded onto a 4-20 % Tris-glycine gel (Bio-Rad). Protein from the gel was transferred to a PVDF membrane. The blot was blocked with 5% (*w/v*) milk in tris-buffered saline with 0.1% (*v/v*) Tween-20 (TBST), and αSynWT/ΔN7::YFP was visualised with a mouse anti-GFP antibody (1:1000) (BioLegend clone B34, 902601). A mouse anti-tubulin antibody (1:5000) (Sigma clone DM1A monoclonal, T9026) was used to detect the tubulin loading control. Both anti-GFP and anti-tubulin were visualised with an anti-mouse horseradish peroxidase-coupled secondary antibody (1:5000) (Cell Signalling Technology, 7076S). Bands were visualised with Supersignal™ West Pico PLUS Chemiluminescent Substrate.

**Confocal imaging and aggregate quantification**

*C. elegans* were bleach synchronised and imaged at day 1, 4 or 8 of adulthood for confocal imaging experiments. Nematodes were immobilised on 2 % (*w/v*) agarose pads with 25 mM sodium azide. The head regions of *C. elegans* were imaged using a Zeiss LSM880 confocal microscope at a magnification of 40 x 1.0 numerical aperture objective. YFP was visualised using an excitation at 514 nm. Z-stacks through the head region were collected for 15-20 worms per experimental condition. Aggregate quantification (from the tip of the nematode to the end of the terminal pharyngeal bulb) was carried out in ImageJ using a partially-automated analysis pipeline, where puncta larger than 1 $\mu m^2$ were considered as aggregates. The results from three biological repeats were collated and error bars shown on graphs are SEM. A two-way ANOVA couple with a post-hoc Šidák multiple comparisons test was carried out in GraphPad Prism 8 to determine statistical significance.

***C. elegans* motility assay**

Age synchronised nematodes were transferred to drop of M9 on an unseeded NGM plate. *C. elegans* thrashing was recorded on a camera attached to the microscope eyepiece for 15 sec each time. A minimum of 20 worms were videoed per condition. The thrashing rates were calculated using the wrMTrck plugin on ImageJ (16). The experiment was repeated three times for each age. A two-way ANOVA couple with a post-hoc multiple comparisons Tukey test was carried out in GraphPad Prism 8 to determine statistical significance.

#### *C. elegans* lifespan assay

*C. elegans* nematodes were transferred to freshly seeded NGM plates at day 1 of adulthood (120-150 animals per condition). Nematodes were scored each day as alive, dead or censored until none were left alive. *C. elegans* were transferred to new seeded NGM plates every other day to maintain a synchronised population and prevent starvation. The experiment was repeated three times and the results plotted using a Python script adapted from Lifelines (17).

### SI Appendix Figures and Figure Legends

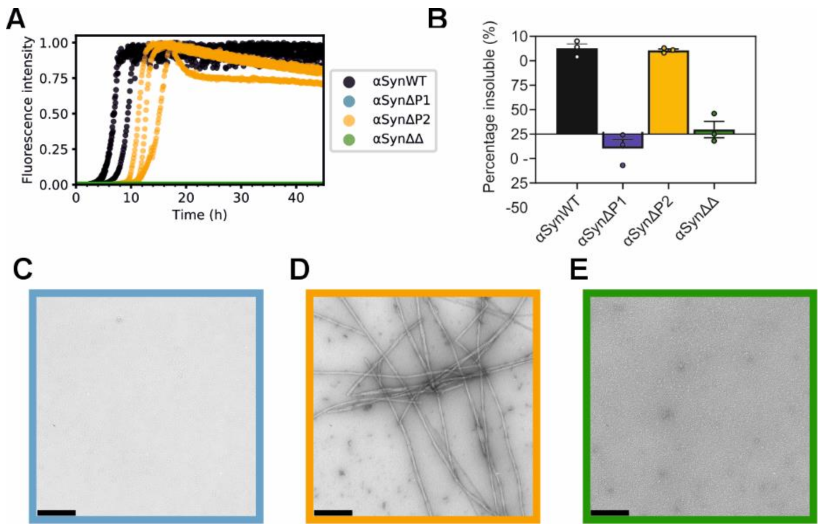


**Fig. S1. Amyloid formation kinetics of the previously-studied deletion variants of αSyn (αSynΔP1, αSynΔP2, and αSynΔΔ). (A)** ThT fluorescence assays with 100 μM αSynWT (black), αSynΔP1 (blue), αSynΔP1 (yellow), and αSynΔΔ (green), in the presence of a Teflon bead (Methods). Conditions are the same as those used for Fig. 1. **(B)** Pelleting assay of the endpoint of the ThT reaction. Note that for **(A)** and **(B)** the data for αSynWT reproduced from Fig. 1 for ease of comparison. **(C-E)** Negative-stain EM micrographs of the endpoint of the ThT assay for **(C)** αSynΔP1, **(D)** αSynΔP2, and **(E)** αSynΔΔ. Scale bar is 250 nm.

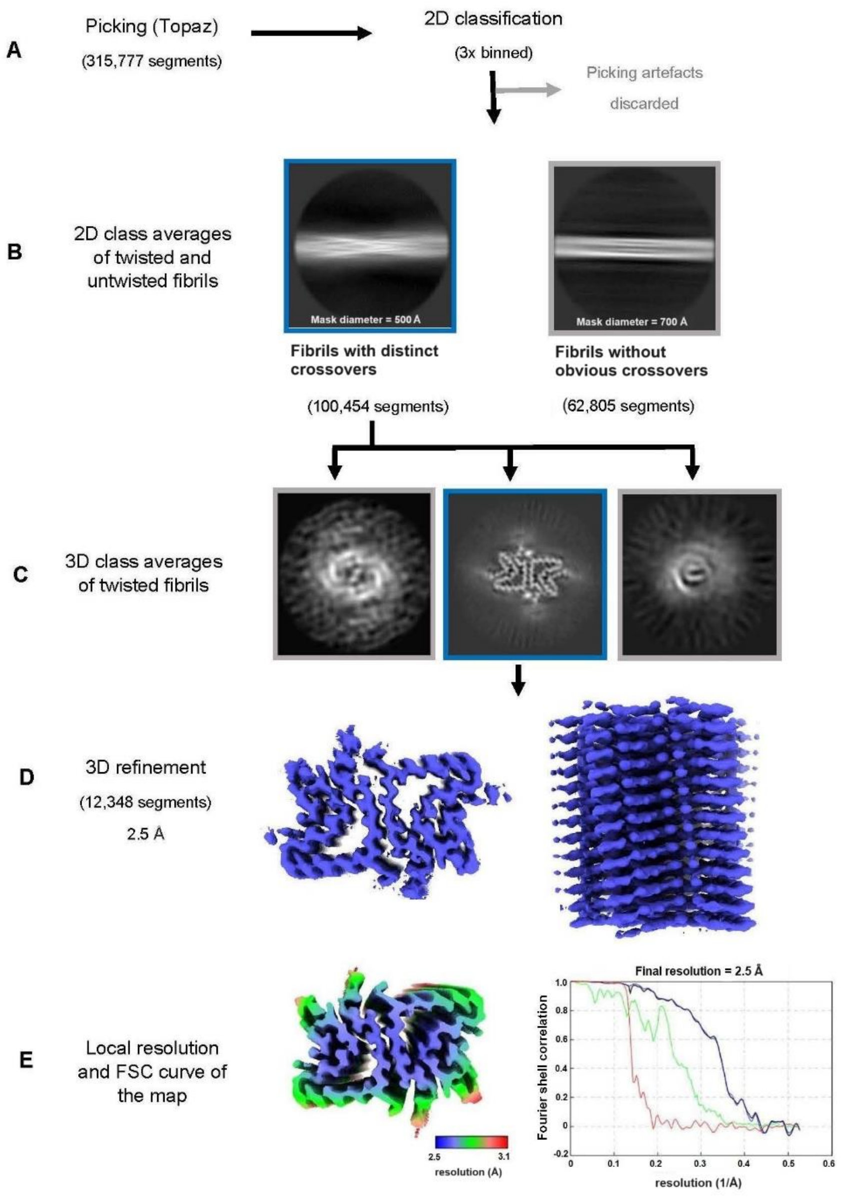


**Fig. S2. Processing flowchart of cryo-EM data analyses. (A)** Topaz picking and initial 2D classification steps to remove picking artefacts. **(B)** Further 2D classifications reveal the presence of two populations of fibrils: one with distinct crossovers, and one without. **(C)** 3D classifications of fibrils with distinct crossover to determine helical rise and twist values. **(D)** Refinement output for αSynΔN7 after Bayesian polishing. **(E)** Local resolution coloured map using RELION4.0 and the FSC curves from the final refinement. The black line shows corrected deposited map, blue shows FSC masked maps, green shows FSC unmasked maps, and red shows FSC phase randomized masked maps values.

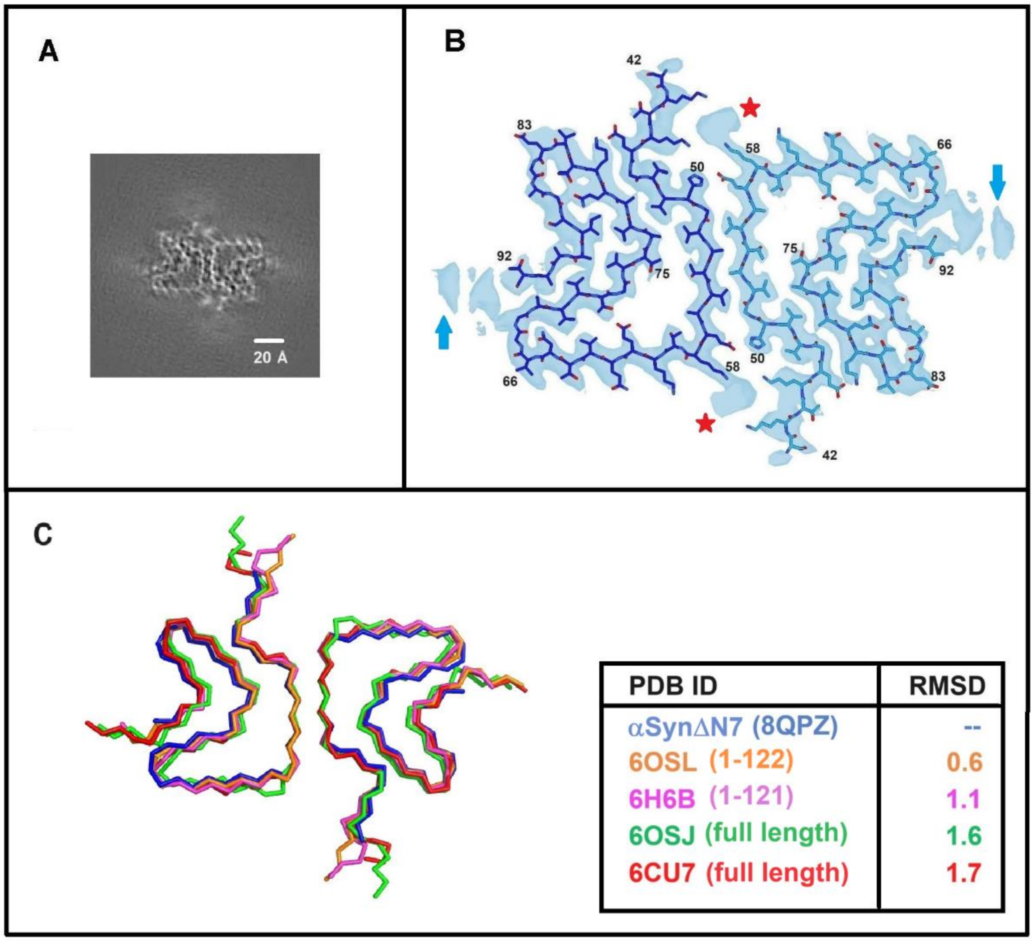


**Fig. S3. αSynΔN7 structure and comparison to other known αSyn structures. (A)** Central slice of the resulting map from 3D classification of αSynΔN7 amyloid fibrils. **(B)** αSynΔN7 fibril structure and associated cryo-EM map. The blue arrows indicate density of residues in the unstructured C-terminal region which could not be modelled into the map. The red stars indicate a non-proteinaceous density. **(C)** Comparison of the αSynΔN7 fibril structure to similar previously solved structures (6, 18) of αSynWT (6OSJ, 6CU7) and its truncated variants (6OSL, 6H6B) show similar filament folds and fibril structures. RMSD values for $C_{\alpha}$ atoms (Å) from the best aligned chains of each structure with αSynΔN7 are shown in the table.

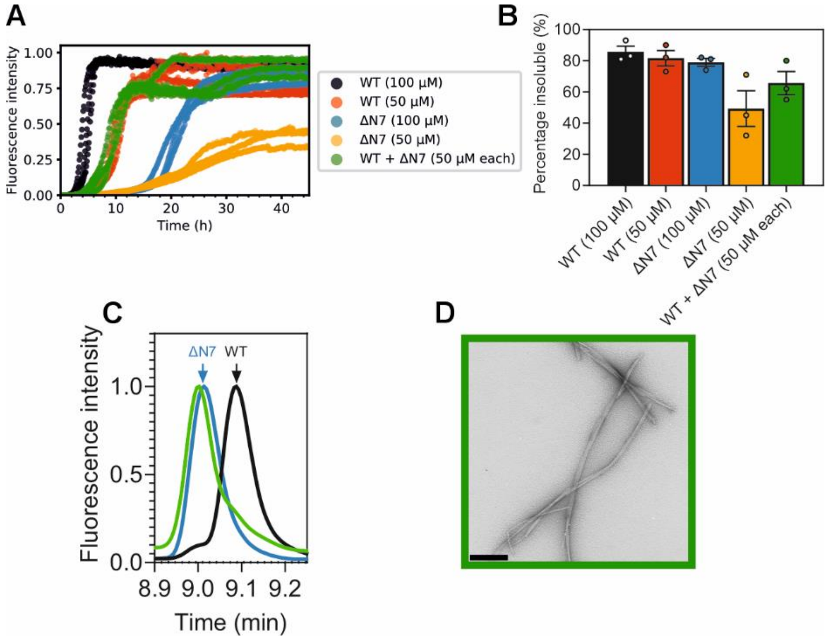


**Fig. S4. Effect of mixing αSynWT and αSynΔN7 monomers on amyloid formation**. **(A)** Fibril formation kinetics of αSynWT and αSynΔN7 when co-incubated in a ThT assay. Controls used are 50 μM and 100 μM of each protein alone. Data are normalised to the maximum ThT signal in the experiment. Three replicates of the experiment are shown in the plot. **(B)** Quantification of the insoluble fraction at the endpoint of ThT assays from three biological repeats, as determined by centrifugation and SDS-PAGE. Error bars are SEM. **(C)** RP-HPLC trace for the soluble fraction of the endpoint of the ThT assay resulting from the co-incubation of αSynWT and αSynΔN7 (green). Profiles of monomeric αSynWT (black) and αSynΔN7 (blue) are displayed to show that the majority of remaining soluble protein at the end of the ThT assay for the co-incubation of 50 μM each of αSynWT and αSynΔN7 is αSynΔN7. Data are normalised to the maximum intensity of each trace. **(D)** Negative-stain EM micrograph of the ThT endpoint after co-incubation. Scale bar is 250 nm.

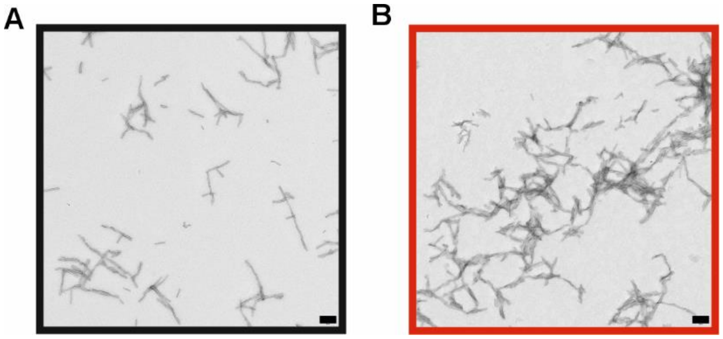


**Fig. S5. Negative-stain TEM images of fibril seeds. (A)** αSynWT and **(B)** αSynΔN7 fibril seeds formed by sonication of the product of fibril growth in 96 well plates. Scale bar is 250 nm.

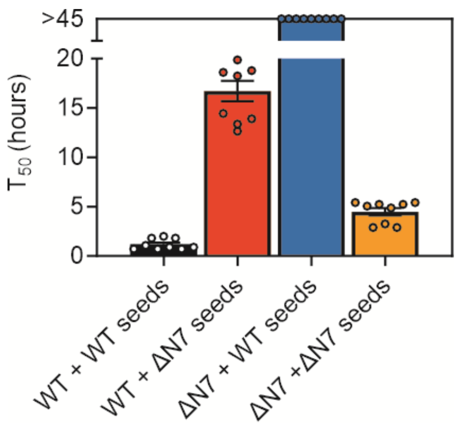


**Fig. S6. $T_{50}$ values for the seeded fibril growth assays shown in Fig. 2**. Nine data points for each reaction were taken from three biological repeats, each containing three technical replicates. For instances when no clear plateau phase was reached, the $T_{50}$ was recorded as >45 h. Error bars are SEM. See also *SI Appendix,* Table S3.

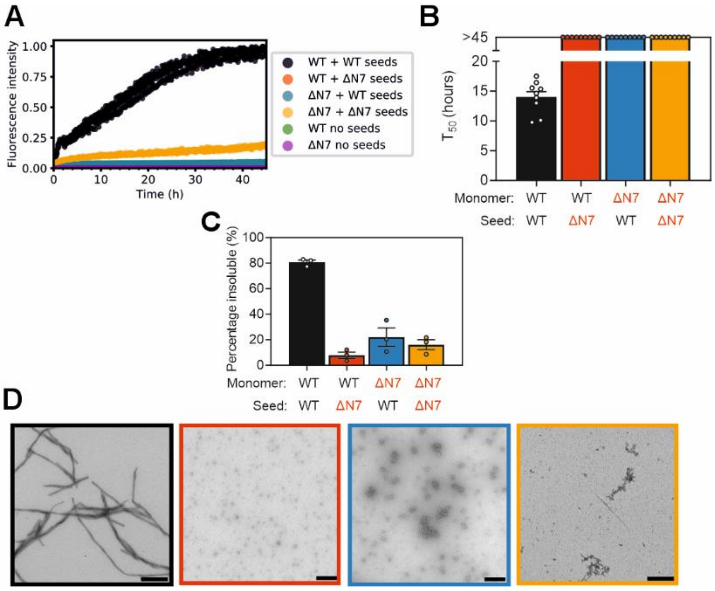


**Fig. S7. Seeded fibril growth kinetics from unsonicated fibril seeds. (A)** Seeded fibril growth for self- and cross-seeding of αSynWT and αSynΔN7 monomers with unsonicated preformed fibril seeds *(*10% *(v/v))*. Data are normalised to the maximum fluorescence of the dataset. Note that for 'WT no seeds' (green) and 'ΔN7 no seeds' (purple) there is no increase in ThT fluorescence signal so the data cannot be seen readily behind each other. **(B)** Corresponding $T_{50}$ values for the unsonicated seeding reactions. For instances when no clear plateau phase was reached, the $T_{50}$ was recorded as >45 h. See also *SI Appendix,* Table S4. **(C)** Quantification of the insoluble fraction at the endpoint of ThT assays. **(D)** Negative-stain electron micrographs of the ThT endpoints, coloured as in **(B)**. Scale bar = 250 nm.

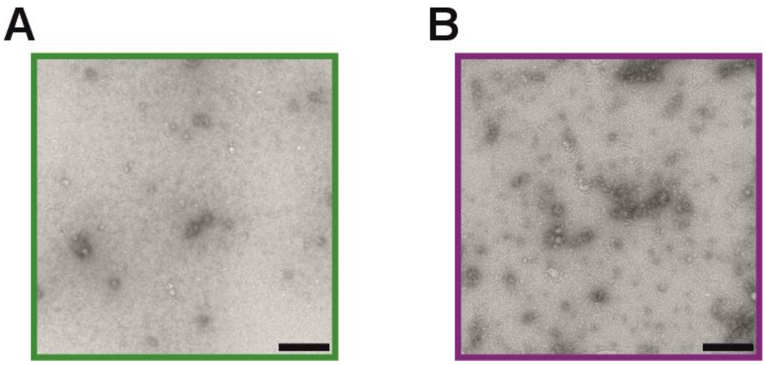


**Fig. S8. Oligomer result from unseeded quiescent incubation of αSynWT and αSynΔN7.** Representative negative stain TEM images of **(A)** αSynWT and **(B)** αSynΔN7 after quiescent incubation in PBS for 45 h at 37 °C. Scale bar = 250 nm.

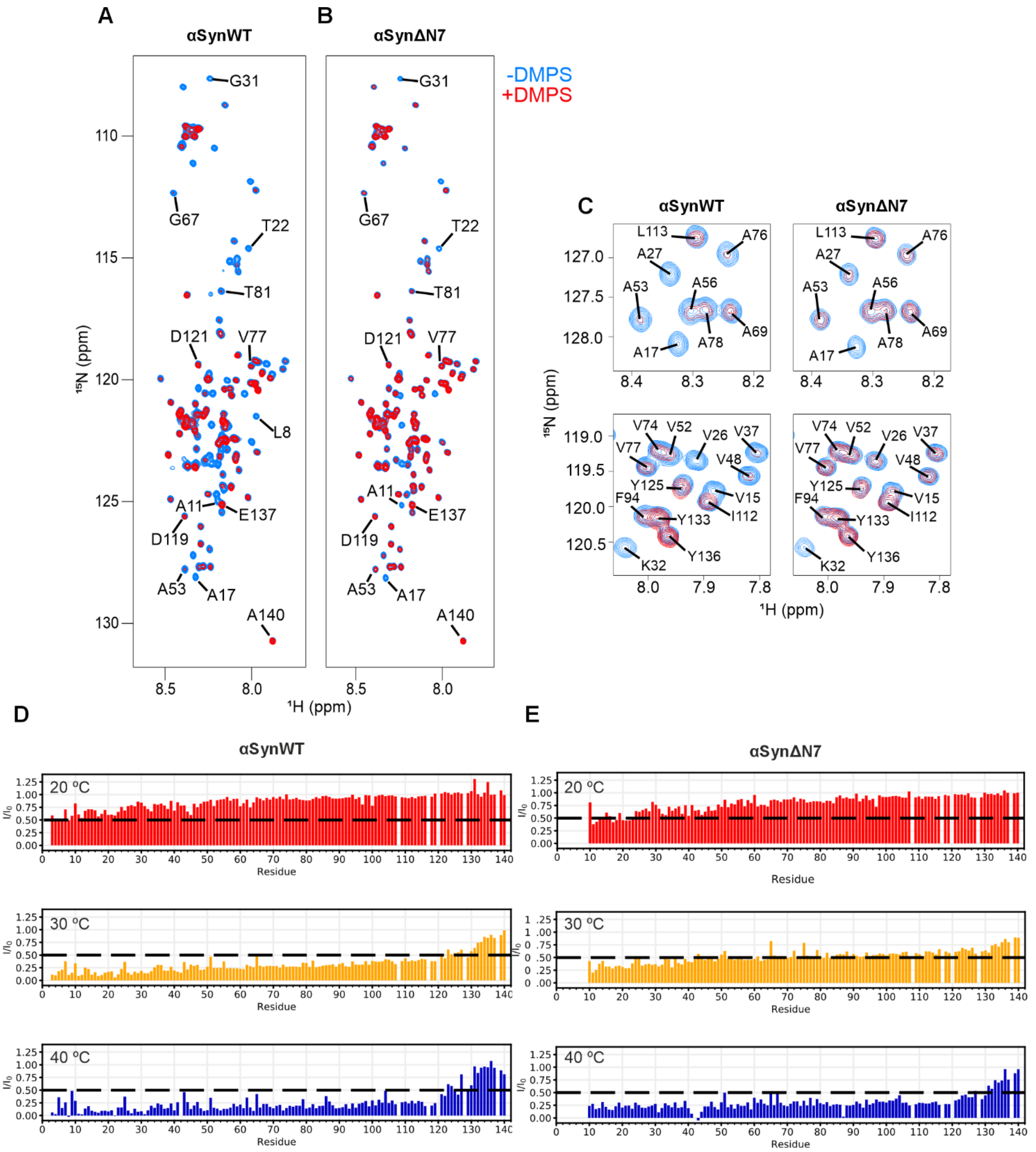


**Fig. S9. $^{1}$H-$^{15}$N HMQC NMR spectra of αSynWT and αSynΔN7 binding to DMPS liposomes at different temperatures. (A, B)** $^{1}$H-$^{15}$N HMQC spectra for αSynWT and αSynΔN7, respectively, in the presence (red) or absence (blue) of DMPS liposomes, collected at 30 °C and an LPR of 8:1. **(C)** Zoom of different regions of the spectra, with individual resonances labelled and coloured as in **(A, B)**. **(D,**

**E)** Intensity ratios at 20 °C (red), 30 °C (yellow), and 40 °C (blue) for αSynWT and αSynΔN7, respectively. The dashed line depicts $I/I_0$ of 0.5. A protein concentration of 25 μM was used and an LPR of 8:1.

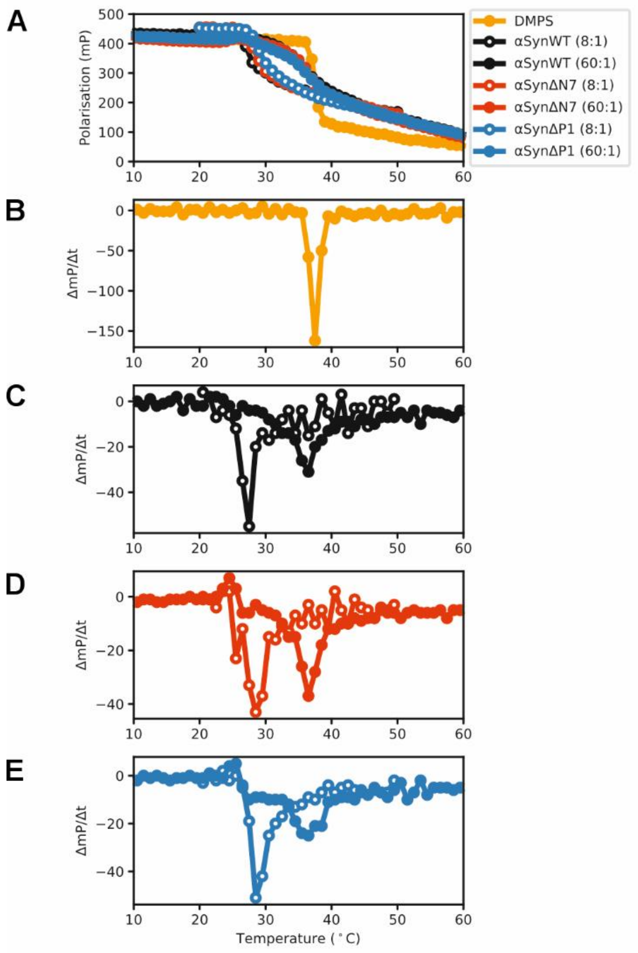


**Fig. S10. Monitoring DMPS membrane fluidity and lipid packing upon incubation with αSynWT and αSynΔN7 monomers.** The experiments were performed using 1,6-diphenylhexa-1,3,5-triene (DPH). **(A)** Change in DPH fluorescence polarisation in DMPS liposomes with temperature. The first derivative of the data in **(A)** is shown for **(B)** DMPS alone or with **(C)** αSynWT, **(D)** αSynΔN7 or **(E)** αSynΔP1 monomers at LPRs of 8:1 and 60:1 (open and closed symbols, respectively). Experimental details are described in *SI Appendix*, Methods.

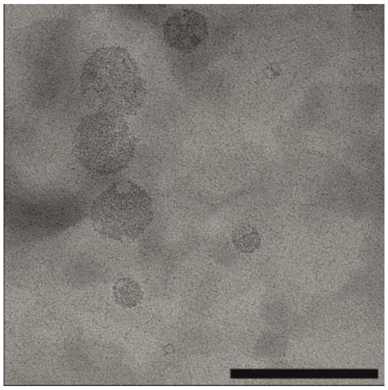

**Fig. S11. Negative-stain image of DMPS liposomes**. Samples were incubated at 30 °C for 45 h in the absence of protein. Scale bar = 250 nm.

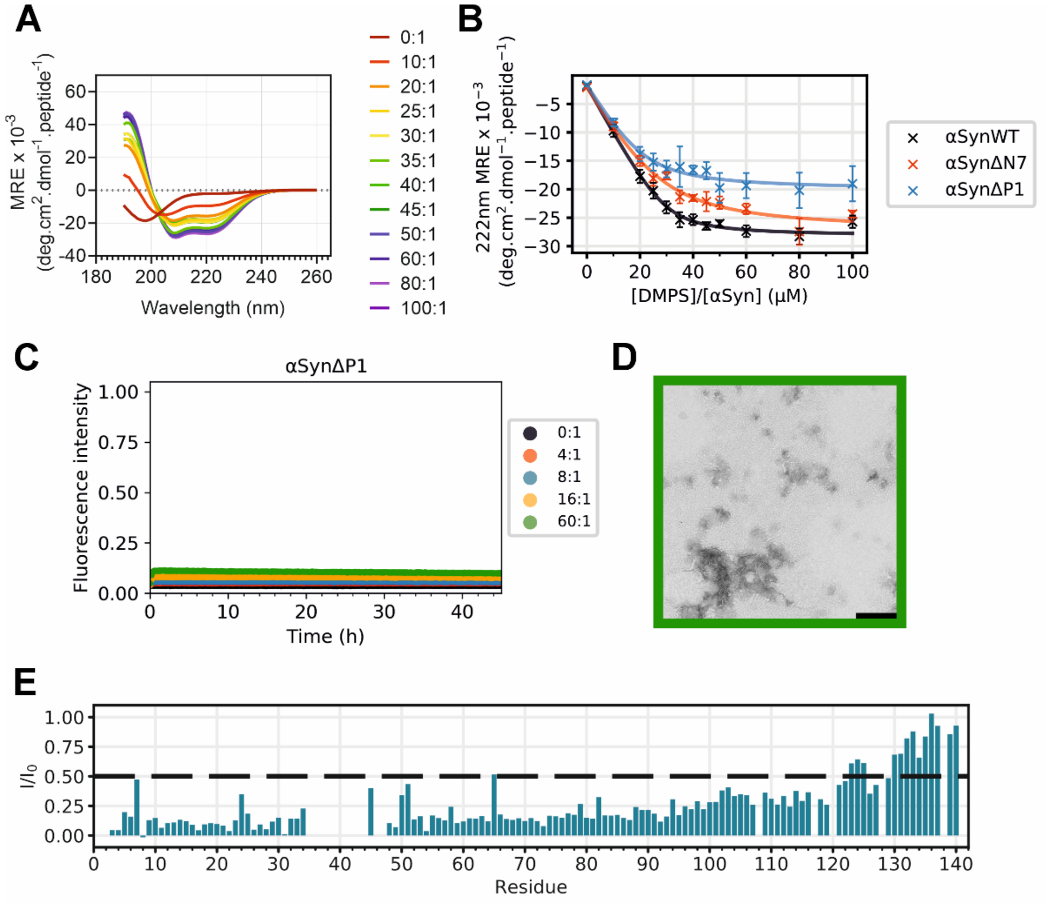


**Fig. S12. Characterisation of the behaviour of αSynΔP1 in the presence of DMPS liposomes. (A)** Far-UV CD spectra of αSynΔP1 at different LPR values as indicated in the key. **(B)** MRE values at 222 nm at different LPR values (blue). Fitting the titration data yielded values of the $K_d$ and L values of 4.2 ± 2.0 μM and 22 ± 4 lipid molecules per molecule of αSynΔP1. Data for αSynΔN7 (red) and αSynWT are reproduced from Fig. 3 for comparison. **(E)** Intensity ratios for each assigned resonance of αSynΔP1 obtained from $^{1}H$-$^{15}N$ HMQC NMR experiments with or without liposomes at a [DMPS]:[αSynΔP1] of 8:1. The dashed line is at $I/I_0$ of 0.5. Note that residues 36-42 are deleted in αSynΔP1. **(C)** ThT assay for αSynΔP1 incubated at different LPRs (indicated in key). Data are normalised to maximum value obtained for αSynWT obtained at an LPR of 16:1 (from Fig. 4A). **(D)** Negative stain electron micrograph of endpoint of incubation of DMPS liposomes with αSynΔP1 at an LPR of 60:1. Scale bar = 250 nm. All experiments were obtained at 30°C.

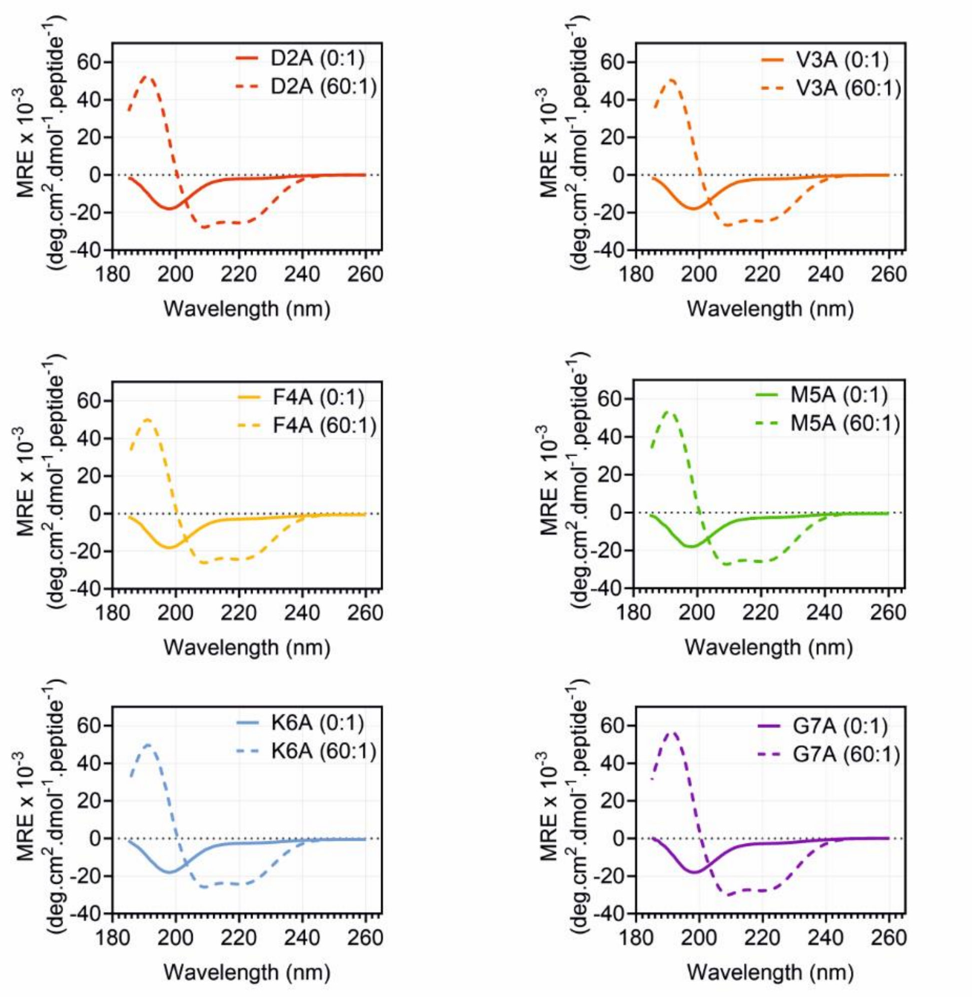


**Fig. S13. Far-UV CD spectra of alanine scan variants of αSyn (residues 2-7).** Spectra were acquired in the absence (solid line) or presence (dashed line) of DMPS liposomes. An LPR of 60:1 was used. All spectra were obtained at 30 °C.

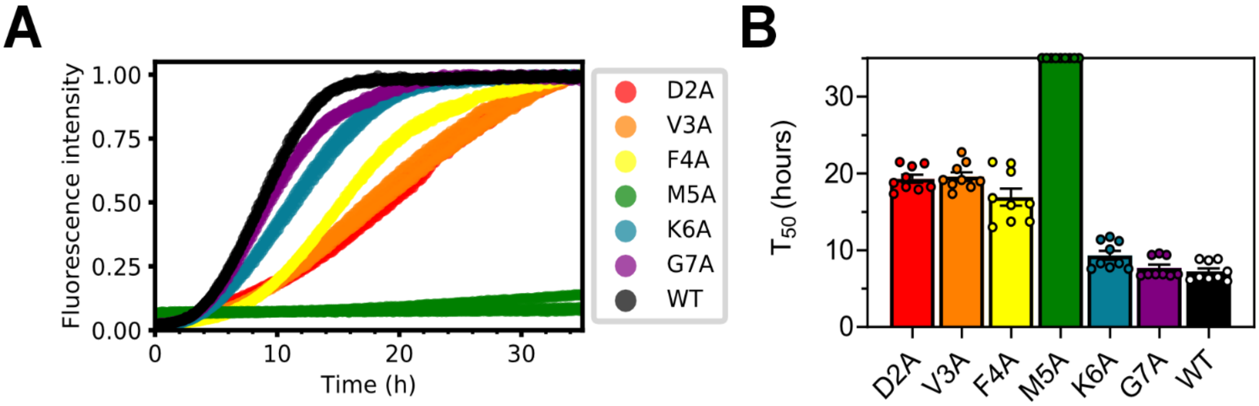


**Fig. S14. Fibril formation kinetics of alanine scan variants of αSyn in the presence of DMPS liposomes. (A)** Fibril formation kinetics for the six alanine scan variants measured by ThT fluorescence. 50 µM protein was incubated in the presence of DMPS liposomes at an LPR of 8:1, 30°C. Three replicates are shown for each variant. **(B)** $T_{50}$ values for each curve of three biological repeats, each containing three replicates (*SI Appendix,* Table S5). Note that for M5A (green) no clear plateau was reached after 35 hours and so the $T_{50}$ could not be determined. Error bars are standard error of the mean (SEM).

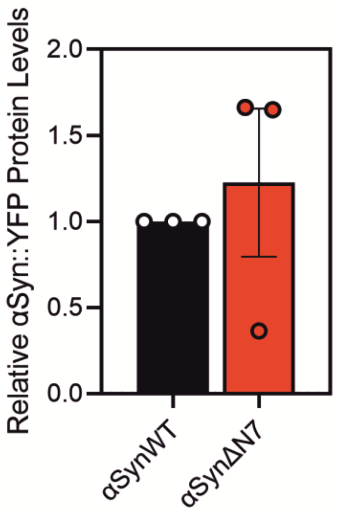


**Fig. S15. Densitometry of the western blots showing protein expression levels of αSynWT::YFP and αSynΔN7::YFP in the body wall muscle cells of *C. elegans***. Expression levels are plotted relative the expression of αSynWT::YFP. Results are shown from three biological repeats and the error bars are SEM.

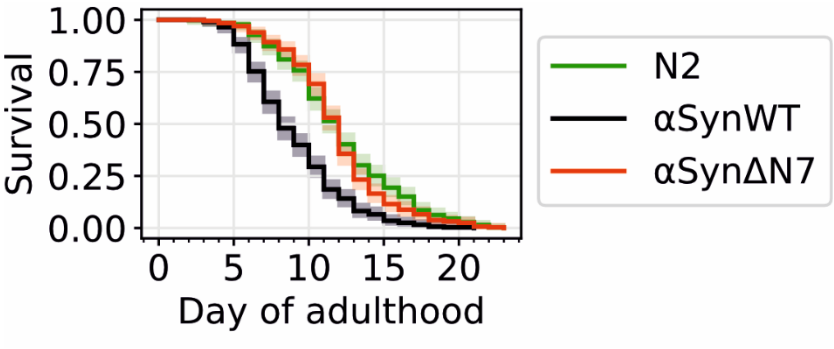


**Fig. S16. Kaplan-Meier plot of the lifespan of C. elegans strains N2 (green), αSynWT (black) and αSynΔN7 (red)**. n = 120-150 animals per biological repeat and the collated results of three biological repeats are plotted. Shaded area shows the 95% confidence intervals. The median lifespan of the strains tested were 15, 11, and 15 days of life for N2, αSynWT and αSynΔN7, respectively. Note that the results are shown as days of adulthood (i.e. not including the 3 days of development to adulthood). Results of the statistical analysis of the lifespans are reported in *SI Appendix*, Table S6.

### SI Appendix Tables and Table Legends

**Table S1. Rate and yield of *de novo* amyloid fibril formation for αSynWT and αSynΔN7 *in vitro*.** The mean $T_{lag}$ and $T_{50}$ were calculated from nine data points (three biological repeats, each containing three technical replicates). The mean percentage insoluble protein was calculated from three repeat experiments. Errors for all values are standard error of the mean. The data were acquired at pH 7.4, in 96 well plates, with agitation in the presence of a Teflon bead at 37 °C.

| | **αSynWT** | **αSynΔN7** |
|---|---|---|
| $T_{lag}$ (hours) | 5.8 ± 0.3 | 18.6 ± 2.4 |
| $T_{50}$ (hours) | 7.2 ± 0.3 | 23.0 ± 2.5 |
| Percentage insoluble (%) | 87 ± 5 | 85 ± 4 |

**Table S2. Cryo-EM data collection, refinement, and validation statistics for the αSynΔN7 dataset**

| | **αSynΔN7 fibril structure<br>(EMDB-18570)<br>(PDB 8QPZ)** |
|---|---|
| **Data collection and processing** | |
| Magnification | 130,000 |
| Voltage (kV) | 300 |
| Detector<br>Energy filter slit width ($e^-$V) | Falcon4-selectris<br>10 |
| Pixel size (Å) | 0.95 |
| Electron exposure ($e^-/Å^2$) | 45 |
| Exposure rate ($e^-$/pixel/s) | 8 |
| Nominal defocus range (μm) | -1.4 to -2.6 |
| Movies collected | 5,464 |
| Initial particle images (no.) | 163,259 |
| Final particle images (no.) | 12,348 |
| Symmetry imposed | C1 |
| Map resolution (Å)<br>FSC threshold | 2.5<br>0.143 |
| Map resolution range (Å) | 2.5 Å – 3.1 Å |
| Helical parameters<br>Helical twist (°)<br>Helical rise (Å)<br>Crossover (nm) | <br>179.43<br>2.44<br>76 |
| **Refinement** | |
| Initial model used (PDB code) | 6OSL |
| Map sharpening *B* factor ($Å^2$) | -26 |
| Model Resolution (Å)<br>FSC threshold | 2.4<br>0.143 |
| Model to map correlation | 0.78 |
| Model composition<br>Non-hydrogen atoms<br>Protein residues total<br>αSyn residues modelled<br>Chains per helical layer<br>Helical layers modelled | <br>4116<br>612<br>42-92<br>2<br>6 |
| *B* factors ($Å^2$)<br>Protein | <br>106 |
| R.m.s. deviations<br>Bond lengths (Å)<br>Bond angles (°) | <br>0.004<br>0.696 |
| Validation<br>MolProbity score<br>Clashscore<br>Poor rotamers (%) | <br>2.2<br>9.3<br>8.6 |
| Ramachandran plot<br>Favored (%)<br>Allowed (%)<br>Disallowed (%) | <br>100.0<br>0.0<br>0.0 |

**Table S3. Half-times ($T_{50}$) and percentage insoluble protein for αSynWT and αSynΔN7 seeded ThT assays containing sonicated amyloid fibril seeds.** Where no clear plateau was reached after 45 hours, the result is indicated by '>45'. See also Fig. 2 and *SI Appendix* Fig. S6.

| | WT + WT seeds | WT + ΔN7 seeds | ΔN7 + WT seeds | ΔN7 + ΔN7 seeds |
|---|---|---|---|---|
| $T_{50}$ (hours) | 1.2 ± 0.2 | 16.7 ± 1.0 | >45 | 4.5 ± 0.4 |
| Percentage insoluble (%) | 87 ± 2 | 75 ± 5 | 28 ± 10 | 70 ± 5 |

**Table S4. Half-times ($T_{50}$) and percentage insoluble protein for αSynWT and αSynΔN7 seeded ThT assays using unsonicated amyloid fibril seeds.** Where no clear plateau was reached after 45 hours, result is indicated by '>45'.

| | **WT + WT seeds** | **WT + ΔN7 seeds** | **ΔN7 + WT seeds** | **ΔN7 + ΔN7 seeds** |
|---|---|---|---|---|
| $T_{50}$ (hours) | 14.0 ± 0.9 | >45 | >45 | >45 |
| Percentage insoluble (%) | 81 ± 2 | 8 ± 2 | 22 ± 7 | 16 ± 4 |

**Table S5. Half-times ($T_{50}$) for alanine scan variants in the lipid-dependent ThT assays.** Where no clear plateau was reached after 35 hours, result is indicated by ‘>35’.

| | D2A | V3A | F4A | M5A | K6A | G7A | WT |
|---|---|---|---|---|---|---|---|
| $T_{50}$ (h) | 19.3 ± 0.5 | 19.6 ± 0.6 | 16.9 ± 1.1 | >35 | 9.3 ± 0.6 | 7.7 ± 0.4 | 7.2 ± 0.4 |

**Table S6. Statistical significance of the difference in lifespan between N2, αSynWT::YFP and αSynΔN7::YFP strains.** A log-rank test followed by a post-hoc Bonferroni correction was used to compare each of the strains. (****) = p < 0.0001. (n.s. = not significant).

| Compared strains | p-value (Bonferroni corrected) |
|---|---|
| N2 vs αSynWT | 1.14E-27 (****) |
| N2 vs αSynΔN7 | 0.621806 (n.s.) |
| αSynWT vs αSynΔN7 | 3.08E-25 (****) |
